# Efficient chemical and enzymatic syntheses of FAD nucleobase analogues and their analysis as enzyme cofactors

**DOI:** 10.1101/2023.01.16.524199

**Authors:** Ateek Shah, Yashwant Kumar, S. Rohan, Amrita B. Hazra

**Affiliations:** Department of Chemistry, Indian Institute of Science Education and Research (IISER) Pune, Dr. Homi Bhabha Road, Pashan, Pune, Maharashtra - 411008, India; Discipline of Chemical Engineering, Indian Institute of Technology Gandhinagar, Palaj, Gujarat - 382355, India

## Abstract

Flavin adenine dinucleotide (FAD), an essential cofactor in cellular metabolism, catalyses a wide range of redox reactions. The organic synthesis of FAD is typically conducted by coupling flavin mononucleotide (FMN) and adenosine monophosphate. The reported synthesis routes have certain limitations such as multiple reaction steps, low yields, and/or difficult-to-obtain starting materials. In this study, we report the synthesis of FAD nucleobase analogues using chemical and enzymatic methods with readily available starting materials achieved in 1-3 steps with moderate yields (10-51%). Further, we demonstrate that *Escherichia coli* glutathione reductase can use these analogues to catalyse the reduction of glutathione. Finally, we show that FAD nucleobase analogues can also be synthesized inside a cell from cellular substrates FMN and nucleoside triphosphates. This lays the foundation for their use in studying the molecular role of FAD in cellular metabolism and as biorthogonal reagents in biotechnology and synthetic biology applications.

## INTRODUCTION

Flavin adenine dinucleotide (FAD) is an essential cofactor in cellular metabolism that assists numerous enzymes in catalysing oxidation-reduction reactions. Many organisms including humans require FAD and produce it from the precursor vitamin B_2_ (riboflavin) via a two-step enzymatic conversion involving two molecules of adenosine triphosphate (ATP). In the first step, the enzyme riboflavin kinase phosphorylates vitamin B_2_ using ATP to form flavin mononucleotide (FMN), which is converted by the enzyme FMN:adenylyltransferase (FMNAT) using another molecule of ATP into FAD (Figure 1A). While the redox capability of FAD is ascribed to the isoalloxazine ring, the ribityl chain, the negatively charged pyrophosphate, and the adenosine groups are implicated in molecular recognition, anchoring, and binding to the enzyme^[1,2]^. Interestingly, the adenosine group is also a recurring moiety in other cofactors such as S-adenosyl methionine (SAM), nicotinamide adenine dinucleotide (NAD^+^), and coenzyme A, and appears to play a similar role in recognition and binding^[2]^.

**Figure 1.**
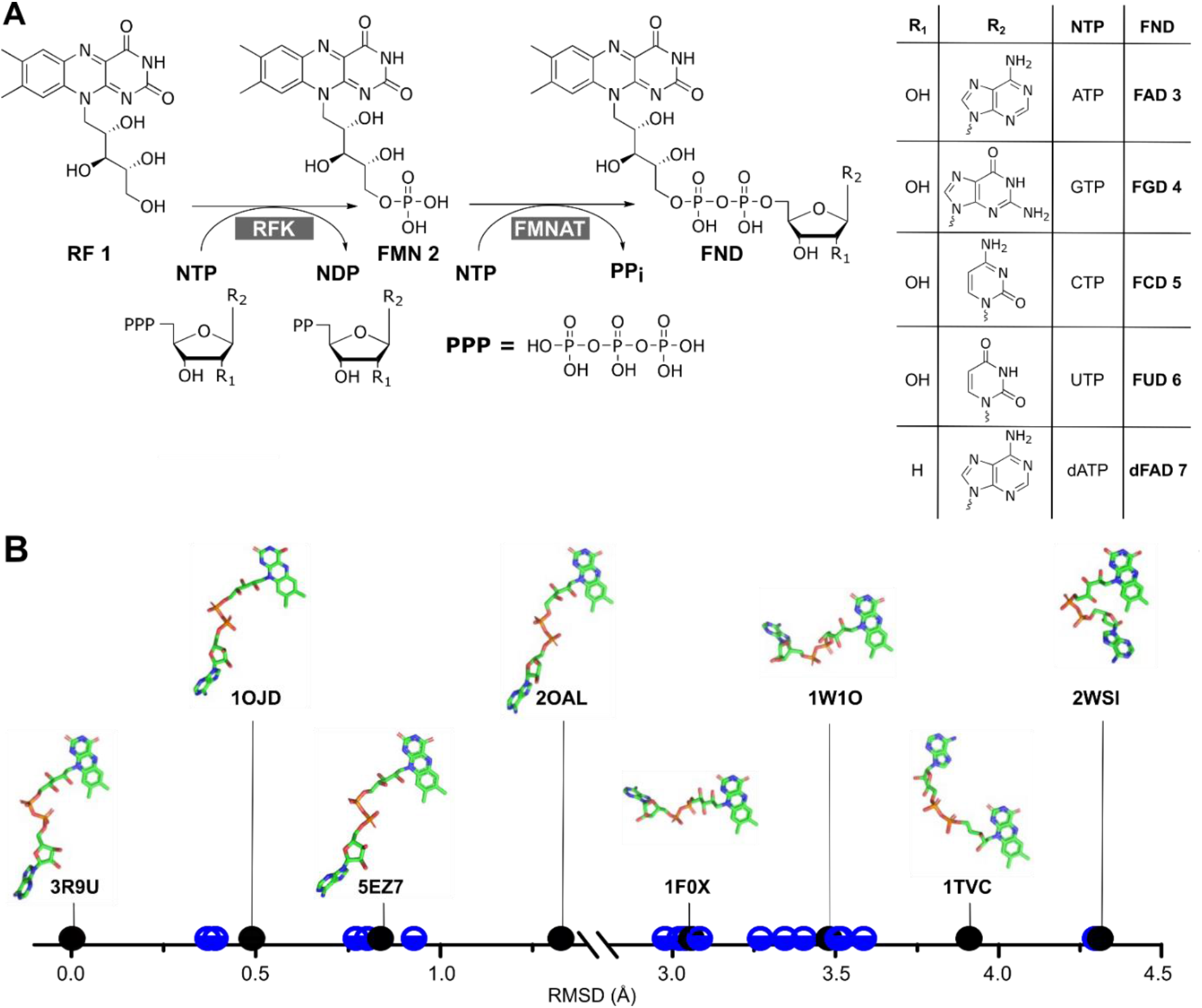
Biosynthesis of flavin adenine dinucleotide (FAD) and its nucleobase analogues and its conformational study in FAD bound proteins. (A) In the cell, riboflavin (RF) is first converted to flavin mononucleotide (FMN) by riboflavin kinase (RFK), followed by adenylation to produce FAD by FMN:adenylyltransferase (FMNAT). If the second step can accommodate other NTPs instead of ATP, then nucleobase analogues of FAD will be formed. (B) The RMSD values were calculated for 24 bound FAD structures from flavoproteins where PDB: 3R9U (Thioredoxin-disulfide reductase from *Campylobacter jejuni*) was taken as reference with 0.0 Å RMSD value (Table S3). Even though the bound FADs in various enzymes show discrete RMSD values, yet each has an extended conformation where the isoalloxazine and adenine ring are mostly apart separated by ~11-16 Å.

Owing to the pivotal role of FAD in cellular metabolism, several analogues have been created for investigating its structural features and as mechanistic probes for studying flavoprotein function. Three major classes of FAD analogues are synthesized – ones with modifications at the (i) isoalloxazine ring, (ii) ribityl chain, and (iii) adenosine moiety. Fluorinated, thiolated, or other isoalloxazine ring variants of FAD are used to examine the environment of the active site of flavin-binding proteins^[3,4]^. Also, antibacterial agents roseoflavin and 5-(3-(4-fluorophenyl) butyl)-7,8-dimethylpyrido[3,4-b]-quinoxaline-1,3(2H,5H)-dione (5-FDQD) display significant pharmacological effects on binding to FMN riboswitches, and several halogenated isoalloxazine ring variants show antimalarial activity as glutathione reductase inhibitors^[5–7]^. Structural variations to the ribityl chain have also yielded interesting insights into FAD binding and catalytic function. Ribityl side chain analogues 2’-deoxyFAD, arabinoFAD, and 2’-F-arabinoFAD show altered catalysis when bound in the active sites of enzymes lipoamide dehydrogenase, glutathione reductase, and mercuric reductase^[8]^. In contrast, FAD nucleoside analogues are not as extensively studied. One photophysical study explores the influence of the nucleobase on the structural and spectroscopic properties of FAD using purine and pyrimidine ribo- and deoxyribonucleoside analogues and concludes that the former exhibits stronger intramolecular complexing^[9]^. A handful of other studies exist that examine the *in vitro* catalytic ability of a variety of analogues modified at the adenine position with D-amino acid oxidase ^[10–14]^. However, the role of nucleobase substitutions in FAD with regard to their biosynthesis or utilization in enzymes and cells is minimally explored.

In this study, we report an efficient synthesis of FAD nucleobase analogues via two methods – a chemical synthesis route and an enzymatic route. Both routes employ ≤3 steps and result in moderate yields (10-50%). We believe that the change of nucleobase will not affect the catalytic ability of the FAD analogues because the adenine and isoalloxazine rings of FAD are typically in the extended orientation, placed ~11-16 Å apart (the distances between terminal “backbone” atoms), when bound to an FAD-utilizing enzyme^[15]^ (Figure 1B). Hence, we tested the binding and activity of *Escherichia coli* glutathione reductase (*Ec*GR) with these analogues, and found that they all bind, and are able to act as reducing agents although with differing efficiencies. Finally, as NTPs and FMN are both naturally occurring substrates in cellular systems, we heterologously expressed the promiscuous *Methanocaldococcus jannaschii* FMN adenylyltransferase (*Mj*FMNAT) and show that FAD nucleobase analogues can also be synthesized inside a cell. Collectively, our analysis puts forth the possibility of using FAD nucleobase analogues to probe the molecular role of FAD in cellular metabolism and in synthetic biology applications involving unnatural vitamins^[16–21]^.

## RESULTS AND DISCUSSION

### Limitations of existing synthesis methods for FAD and its analogues

The first chemical synthesis of FAD involved the condensation of an adenosine-5’-benzyl phosphorochloridate derivative with various metal salts of FMN, followed by the removal of protective groups^[22]^. This method was slow with multiple steps and yielded very less product (~6%). As an improvement, a one-step condensation of adenosine monophosphate (AMP) with salts of FMN in the presence of trifluoroacetic acid anhydride (TFAA) as a condensing agent was devised^[23]^. This strategy, used for the synthesis of FAD and its nucleobase analogues, has extremely poor yields in the range of 0.5-1% owing to the self-condensation side reactions of the substrates^[10]^. Yet, being a one-step process with readily available starting materials, it has been used in a few instances to create nucleobase cofactor analogues^[24,25]^. In a slightly modified approach, carbodiimide is used as condensing agent to synthesise FAD from FMN and AMP, however, yields remained poor^[11]^. An approach to improve the yield involves activation of one of the phosphate substrates as a phosphoramidate or phosphoromorpholidate followed by attack by the other phosphate^[9,12,26,27]^. This route improves the yield to the range of 25-40% but the low reactivity of the phosphoramidate or phosphoromorpholidate results in the reaction time extending to 4-7 days. When diphenyl phosphorochloridate is used as an activating agent for FAD synthesis, the yield is 70-80% and the reaction is complete within a few hours^[28]^. However, when FAD and NAD analogues were synthesized by this method, the yield drops to 25-30%^[13,14,29]^ Another method uses riboflavin phosphorothioate with silver salt of adenosine monophosphate, achieving 51% yield in 5 hours^[30]^. However, obtaining of the starting material is a limiting factor, as it is not commercially available. Each of these methods to chemically synthesize FAD nucleobase analogues are limited by yield, number of steps, time of synthesis, or ease of procuring the starting substrates (summarized in Table S1). The bifunctional FAD synthetase from *Corynebacterium ammoniagenes* has been systematically standardized for the enzymatic synthesis of FAD analogues modified at the isoalloxazine ring and the ribityl chain, however, this method does not work for FAD nucleobase analogues^[31,32]^. The FAD synthetase enzyme has two active sites – one site for phosphorylation of the riboflavin to make FMN which accommodates a variety of modified isoalloxazine and ribityl chain analogues and the second site for combining FMN and AMP to produce FAD. As the active site is very specific for the nucleotide substrate, FAD nucleobase analogues cannot be synthesized^[33]^. Owing to all these limitations, obtaining sufficient quantities of these nucleobase analogues for conducting rigorous chemical characterization remains to be done, and their use in biological applications has been minimal.

### An efficient method for the organic synthesis of FAD and its analogues

To develop a robust synthesis method for FAD nucleobase analogues, we set out to explore the direct condensation route of FMN and a nucleoside monophosphate (NMP) to produce FND using alternate activating agents. Even though direct condensation routes have historically produced very low yields, they offered advantages such as (i) lesser number of steps, (ii) short reaction time, and (ii) ease of accessing the starting materials. To increase the efficiency of the condensation reaction, we chose activating agent N-methylimidazole based on the precedence of a reported synthesis of deoxynucleoside 5’-triphosphates (dNTP) ^[34]^. This was achieved by acylation of the hydroxyl groups on the phosphate of NMP with TFAA followed by reaction with N-methylimidazole to form the corresponding dNMP-N-methylimidazolide and finally, coupling with inorganic pyrophosphate to produce the dNTP with 89-92% yield. Since then, this strategy has been used for the synthesis of UDP-α-D-galactofuranose, other sugar nucleotides and di/triphosphate-containing metabolites ^[35–38]^. In this study, we adapted this method to create FAD nucleobase analogues and have achieved a 3-step reaction (~20% yield) or 2-step reaction (~10% yield), both conducted in less than 5 hours.

### Synthesis of acetylated FMN (FMNAc)

The starting substrates for the synthesis of FAD nucleobase analogues by this method are FMN **2** and NMP **9**. As FMN is sparingly soluble in organic solvents, we acetylated FMN with acetic acid anhydride in the presence of perchloric acid to form tri-acetylated FMN (FMNAc **8**)^[22,39]^ (Figure 2). The reaction yielded a mixture of acetylated FMN products (FMN acetylated at one, two or all three hydroxy positions) along with acetylated riboflavin (as riboflavin is present along with FMN **2** in the starting material) (Figure S1). After conducting a workup with chloroform to remove the acetylated riboflavin, FMNAc **8** was purified via C18 reversed-phase chromatography (Figure 3).

**Figure 2:**
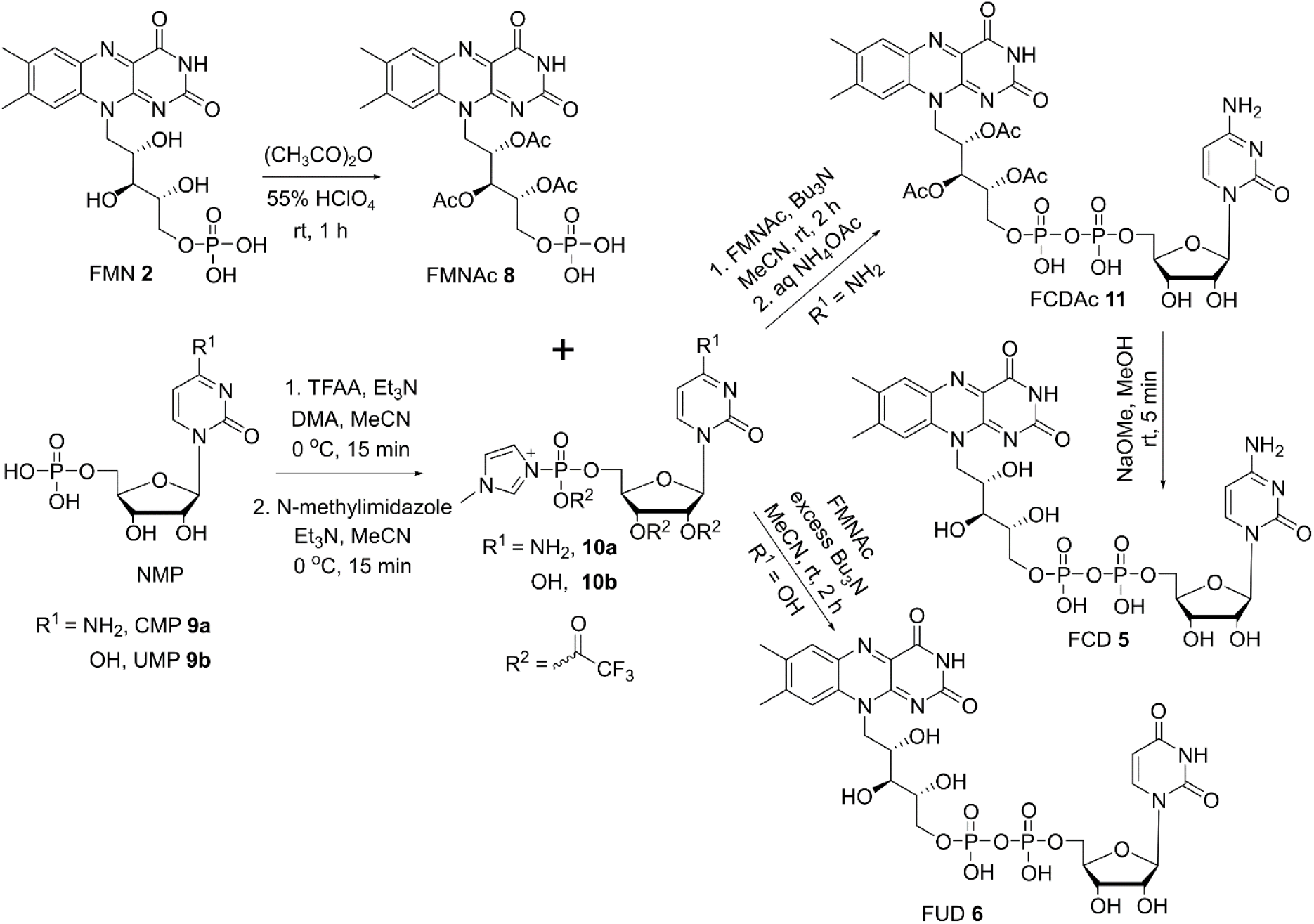
Scheme for the synthesis of FAD analogues FCD and FUD. NMP (CMP/UMP) is first activated by trifluoro acetic anhydride (TFAA), followed by N-methylimidazole, which is then reacted with acetyl-protected FMN (FMNAc) to produce the corresponding acetylated FND (FNDAc). Next, FNDAc can be deacetylated by sodium methoxide to form FND. Alternately, excess tributylamine can be used to skip the step of FNDAc formation (done in the case of FUD synthesis), which reduces the separate deacetylation step, however, it is less efficient, giving a lower yield.

**Figure 3.**
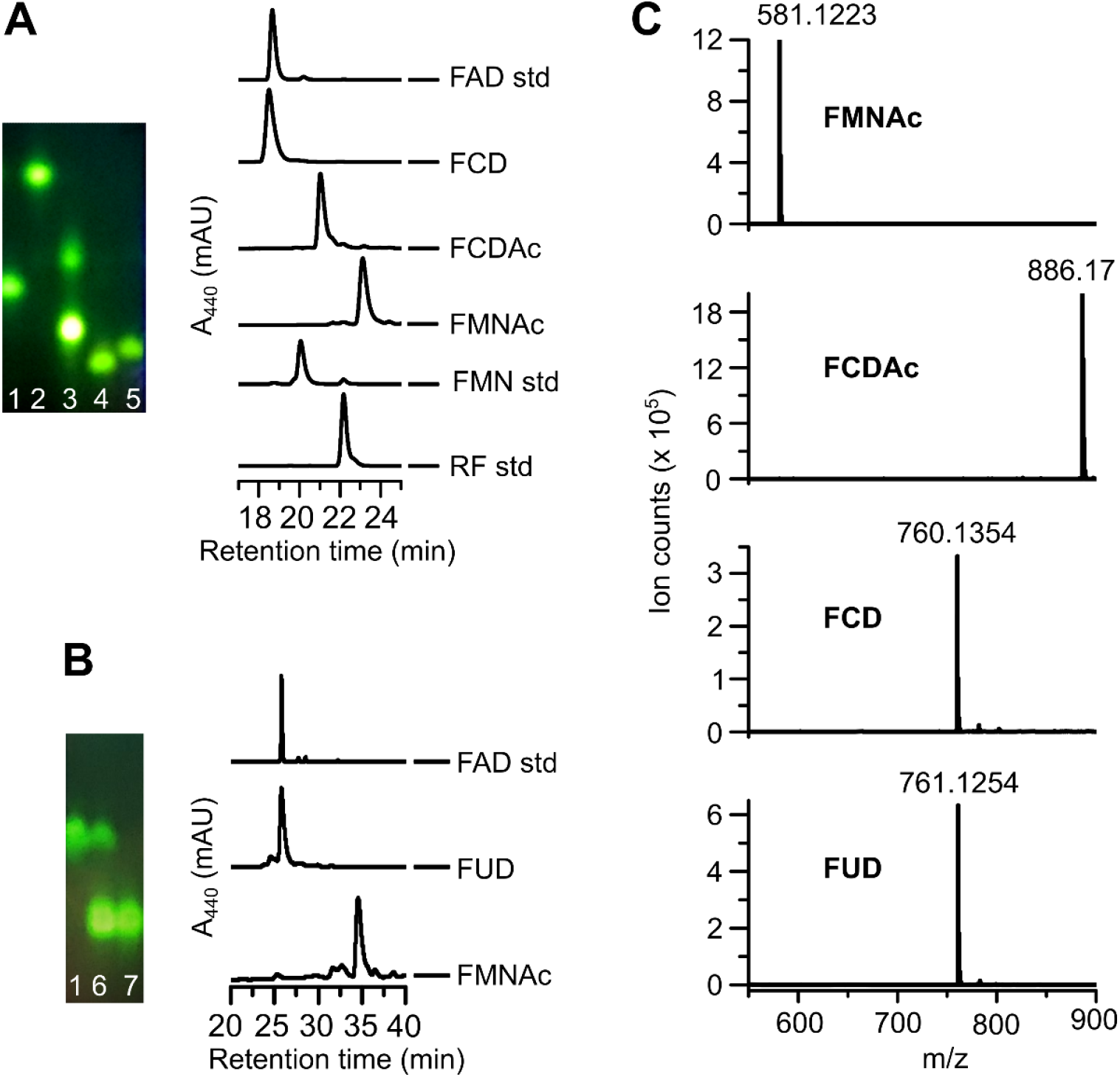
Characterisation of FCD, FUD and their intermediates via. (A) TLC (lane 1-FMN, 2-RF, 3-FMNAc (upper) and FCDAc (lower), 4-FCD, 5-FAD) and HPLC chromatogram showing different retention time of starting material, purified intermediates, and final product for FCD synthesis. (B) TLC (lane 1-FMN, 6-FMN and FUD, 7-FUD) and HPLC chromatogram showing the starting material and purified product for FUD synthesis. (C) Liquid chromatography electrospray ionization mass spectroscopy (LC-ES-MS) negative mode (M-H^-^) of purified intermediates and products of FND synthesis.

### Coupling of FMNAc with CMP or UMP to yield FCD or FUD

To conduct an efficient coupling of FMNAc with CMP **9a**, the phosphate of **9a** was activated with TFAA and further activated by N-methylimidazole to make the N-methylimidazolyl compound **10a** (Figure 2). Finally, **10a** was coupled with FMNAc **8** in the presence of tributylamine. C18 reversed-phase purification yielded ~31% of the product FCDAc **11**. To obtain FCD **5** from FCDAc **11**, an acid or base-catalysed deacetylation is required. However, FCD **5** is not stable under these conditions. Thus, deacetylation of FCDAc **11** was attempted in 50 μM of sodium methoxide in MeOH for short period of time (3-5 min) and the reaction was finally quenched with equimolar HCl solution. Since FCD **5** is insoluble in MeOH, it precipitated out along with other FMN derivatives formed during the course of the reaction. FCD **5** was then purified by reversed-phase chromatography and confirmed by fluorescence thin-layer chromatography (TLC) and high-performance liquid chromatography (HPLC) (Figure 3A, B). The intermediate product FCDAc **11** and final product FCD **5** were further confirmed by mass spectrometry and NMR, where a final yield of 17% was obtained (Figure 3C, 4). The robustness and versatility of this method was verified by the synthesis of FAD **3** via FADAc (figure S7-9). This method is an improvement over previously reported techniques with regard to the number of reaction/purification steps and ease of obtaining starting materials from commercial sources, even though the yield is modest. Also, this method is easily amenable to switching of the nucleotides to synthesize a host of nucleobase analogues of FAD.

To further decrease the number of steps, instead of purifying and then deacetylating the FNDAc, we attempted to conduct the deacetylation after coupling of the monophosphates (Figure 2). The synthesis of FUD **6** was done in the same manner as FCD **5** until the activation of UMP with N-methylimidazole following which we added an excess of tributylamine and triethylamine in the coupling step, which deacetylated FUDAc and FMNAc **8** to FUD **6** and FMN **2,** respectively. FUD **6** was purified and confirmed by LC-MS and NMR (Figures S16-18). This requires two steps as compared to the previous method which needed three steps. This is a significant improvement with regard to time and resources for synthesis and purification of FUD **6**, however, the yield decreases to 11%.

Finally, as the coupling occurs between the terminal phosphate of FMNAc with the activated NMP, in principle, all forms of acetylated FMN (acetylated at one, two or all three hydroxy positions) will yield a mixture of acetylated FND molecules. Thus, the reaction with the acetylated FMN mixture followed by deacetylation should yield FND and the by-product FMN (Figure S1). However, we observed that the FNDAc deacetylation step which utilizes sodium methoxide causes a large loss of yield owing to the formation of a degradation product which resembles riboflavin cyclic-4’,5’-phosphate (cyclic FMN or cFMN)^[40]^. Thus, we conclude that the purification of the triacetylated FMN is an important step in obtaining higher yields of the final product.

### Enzymatic synthesis of FGD and dFAD

Another efficient approach to synthesize FAD nucleobase analogues is tapping into enzymes in the FAD biosynthesis pathway. *In vitro*, *Methanocaldococcus jannaschii* FMN adenylyl transferase (*Mj*FMNAT) was reported to be promiscuous to CTP forming FCD **5** which was identified by LC-ESI-MS using chemically synthesized FCD **5** as a reference^[24]^.

We analysed reactions of *Mj*FMNAT with all the nucleotides and found that *Mj*FMNAT is not only promiscuous for CTP but also for other nucleotides including deoxynucleotides (Figure S2). Hence, we standardized conditions for an efficient *Mj*FMNAT-catalysed enzymatic method for the synthesis of FAD analogues. FGD **4** and dFAD **7** were synthesized by this method, and their yields were quantified (Figure 5). Heterologous overexpression and purification of *Mj*FMNAT was conducted, followed by reaction with FMN and GTP or dATP. We obtained ~90% conversion to FGD **4** and dFAD **7** from FMN **2** and NTP (Figure 6). After HPLC purification of FGD **4** and dFAD **7**, the recovered yield was 51% and 45% respectively.

**Figure 4.**
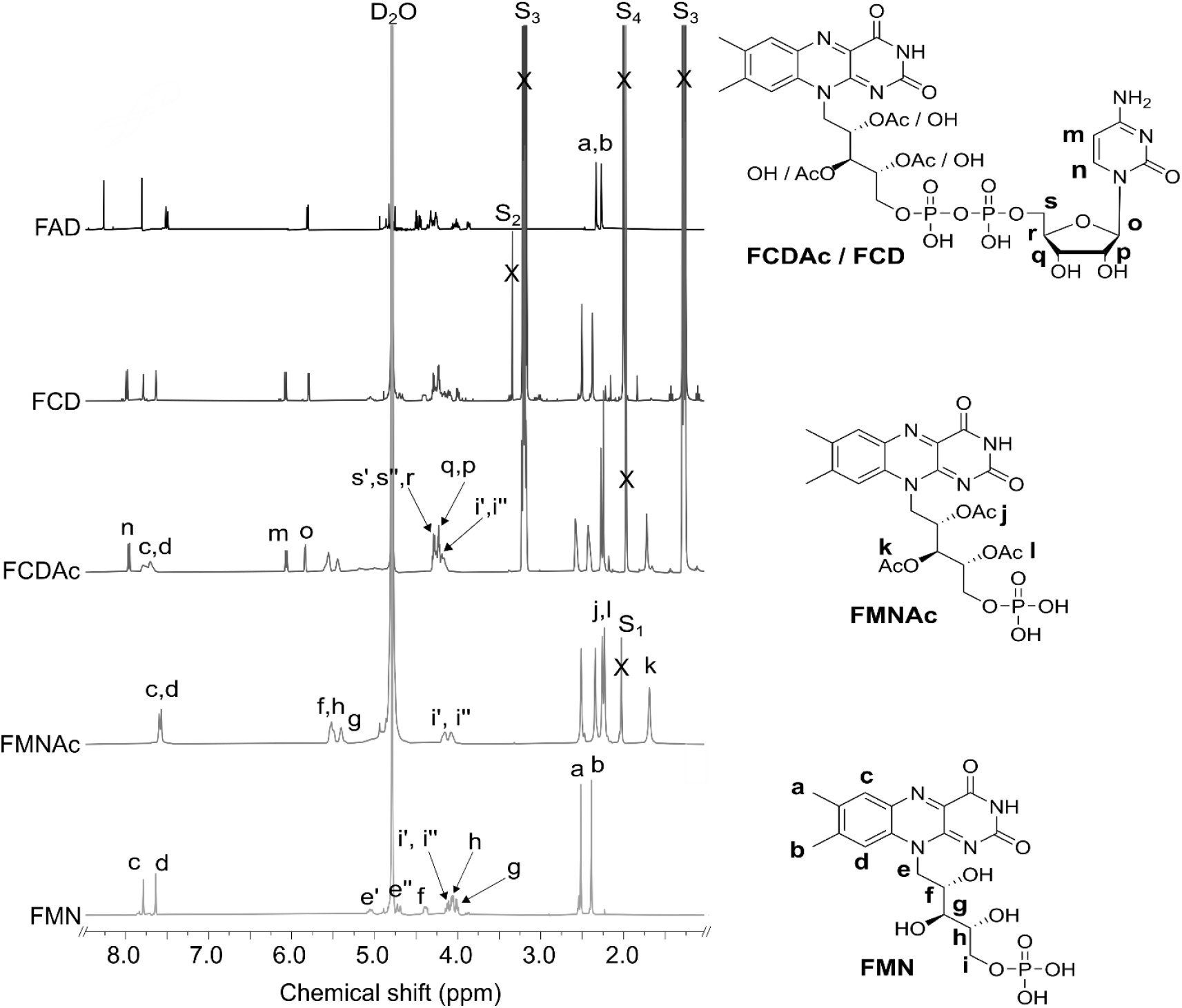
NMR comparison of starting material, intermediates, and products for the synthesis of FCD. FMNAc shows three extra acetyl peaks (j, k, and l), and corresponding CH proton peaks (f, g, and h) shift downfield compared to FMN. FCDAc has additional peaks of the adenosine moiety where two aromatic peaks (m, n) are noteworthy. FCD does not have the three acetyl peaks (no j, k, and l peak) and hence corresponding CH proton peaks (f, g, h)) shift back to the upfield position as in FMN. Peaks S_1_-S_4_ are residual acetone (CH_3_), methanol (CH_3_), triethylamine (CH_2_ and CH_3_), and acetate (CH_3_) solvent peaks respectively. The ‘ and ‘’ symbol on protons at same position represents the different shielding due to adjacent chiral carbon.

**Figure 5.**
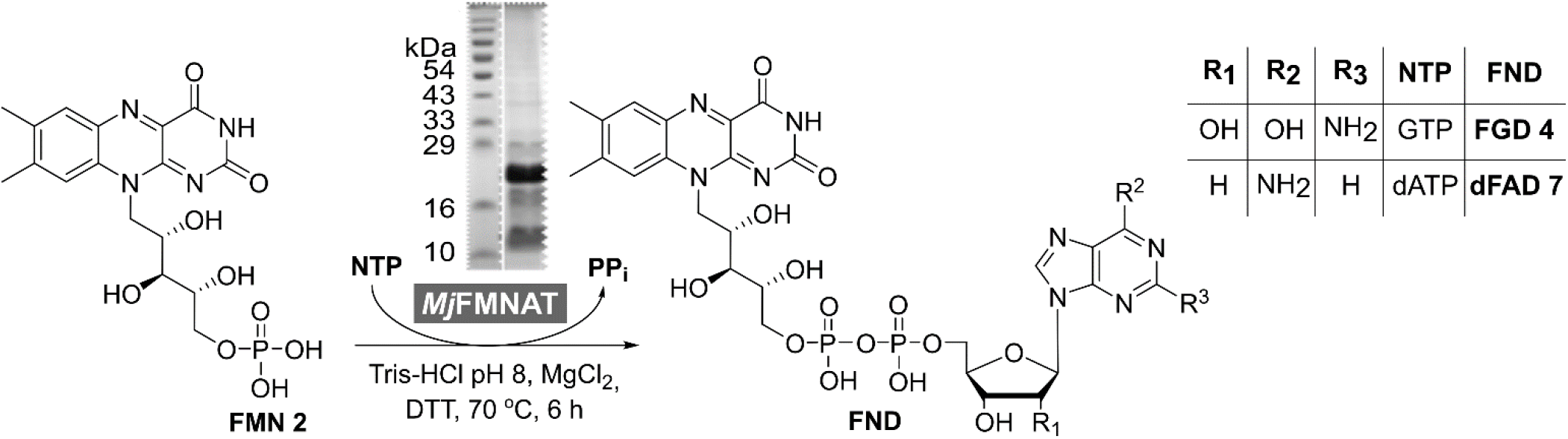
Scheme for the chemo-enzymatic synthesis of FGD and dFAD. *Mj*FMNAT is a promiscuous enzyme for other NTPs which synthesizes the nucleoside analogues of FAD using FMN and NTP.

**Figure 6.**
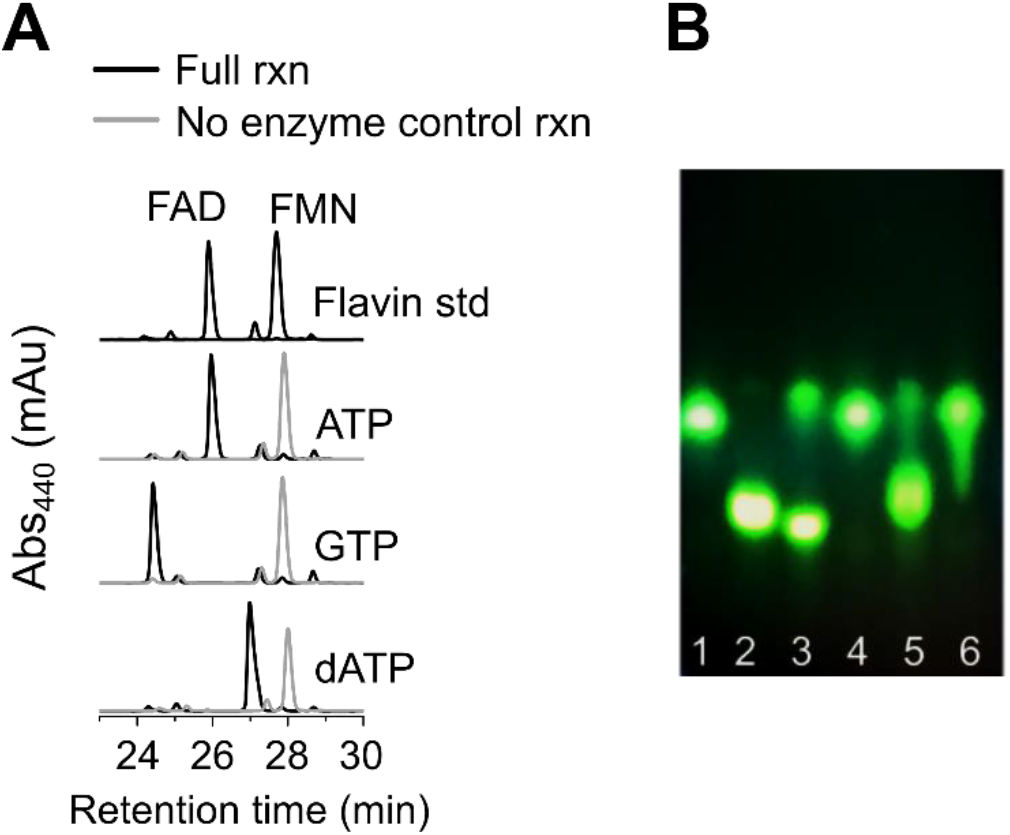
Characterisation of products FAD, FGD, and dFAD using the enzyme *Mj*FMNAT and other NTPs. (A) HPLC chromatogram showing the no enzyme control (gray) overlayed with the full reaction (black). We observed the disappearance of the FMN peak (28 min) in full reaction and appearance of the FND product peak (between 24-27.5 min). (B) TLC. Lane 1-FMN standard, 2-FAD standard, 3-full reaction with GTP, 4-no enzyme control reaction with GTP, 5-full reaction with dATP, and 6-no enzyme control reaction with dATP.

Up until now, the only enzymatic syntheses of FAD analogues have been conducted with *C. ammoniagenes* FADS, however no nucleobase analogues could be synthesized as the enzyme was specific for its requirement for ATP. In this study, we report the first enzymatic synthesis and characterization of a host of FAD nucleobase analogues with moderate yields using the *Mj*FMNAT enzyme.

### UV-visible and fluorescence spectra of FAD analogues

We observed that the UV-vis spectra of FAD analogues are very similar to one another, however, the intensity ratios of λ_270_: λ_370_ and λ_270_: λ_450_ varies (Figure S3A). With respect to FAD, dFAD shows an identical spectrum while FCD, FGD, and FUD show different peak ratios. Also, RF and FMN have low peak ratios compared to FAD and FAD analogues (Figure S3A table). These characteristic ratios may be used to specifically identify FAD nucleobase analogues. The fluorescence spectra of the FAD analogues, we synthesized, are similar to FAD, with small shifts visible at the λ_ex_ and λ_em_ maxima. FAD has the λ_ex_ of 450 nm while FGD and FCD show 445 nm and 452 nm, respectively (Figure S3B). Also, FAD λ_em_ is 525 nm while FGD and FCD display λ_em_ at 523 nm and 525 nm, respectively.

### Utilization of FAD nucleobase analogues as cofactors

Our next goal was to test whether these FAD analogues can be utilized as redox cofactors in enzymes. We chose *E. coli* glutathione reductase (*Ec*GR) as our enzyme of choice. Glutathione reductase uses FADH_2_ to reduce glutathione to its monomeric form and generates FAD (Figure 7A). In the assay, the FAD is converted to FADH2 in the enzyme by NADPH which forms NADP, and the change in absorbance at 340 nm, a result of NADPH oxidation, is used to monitor this reaction^[41]^. We used this method to detect whether *Ec*GR could use the FAD nucleobase analogues we have synthesized to catalyse its reaction. As *Ec*GR elutes with FAD tightly bound to it as prosthetic group, we precipitated the enzyme with saturated ammonium sulphate and reconstituted it with the various FAD analogues. We confirmed the binding of FAD analogues by comparing their absorbance spectra with unbound analogues^[42,43]^. Enzyme bound with FAD, or an FAD analogue shows a red shift when compared to the unbound analogues for 440 nm peak (Figure 7B). Finally, when *Ec*GR reconstituted with FADH_2_ or FADH_2_ analogues were assayed for activity, we found that FAD and its nucleobase analogues are all capable of reducing glutathione (Figure 7C). However, the enzyme is around 50% active with FCD and FUD (0.032 and 0.034 μmoles/min reduction of GSSG by 1 μM enzyme) as compared to FAD (0.06 μmoles/min), even though the enzyme is 80-90% reconstituted with the cofactor (Table S2).

**Figure 7:**
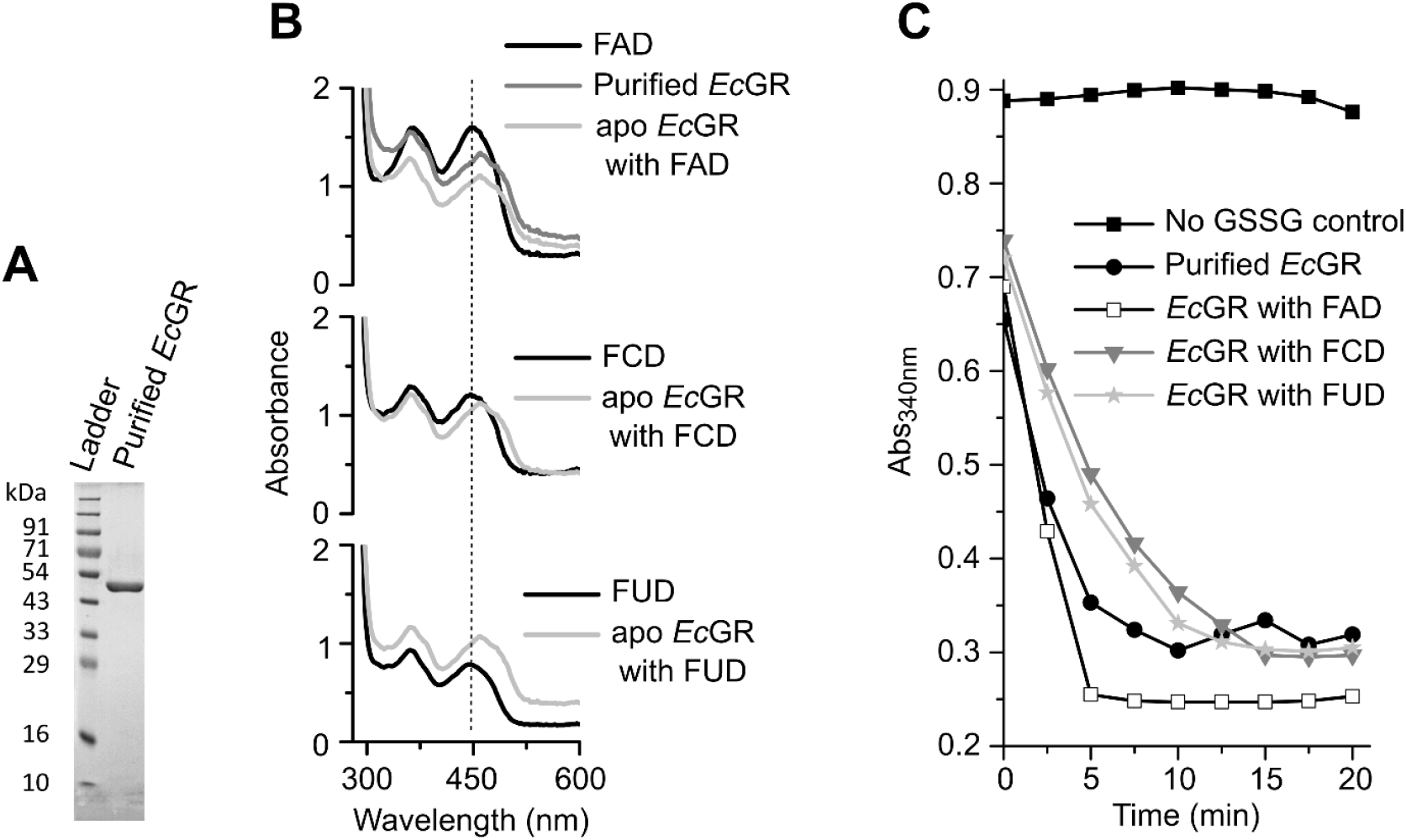
Purification, reconstitution, and characterisation of *Escherichia coli* glutathione reductase (*Ec*GR) with FAD analogues. (A) SDS-PAGE gel showing the purified protein (molecular weight 51 kDa) (B) The absorbance peak of FND at 450 nm undergoes a red shift when apo *Ec*GR is present which indicates that FAD and its analogues are bound to the enzyme. (C) *Ec*GR binds the FAD and its nucleobase analogues and catalyses the reduction of the oxidised glutathione to its monomeric form. The oxidised FND is again reduced by NADPH and the drop in absorbance at 340nm indicates the formation of corresponding NADP.

### Heterologous formation of FAD analogues in *E. coli*

Finally, as our goal is to use these FAD analogues as cofactors in cells, the feasibility of producing them within a cellular system was to be established. To do so, we heterologously expressed the *Mj*FMNAT-containing plasmid in *E. coli*. As *E. coli* already contains *ribCF* which codes for FAD synthetase and is an essential gene that cannot be deleted, we included controls of an empty plasmid and that of a plasmid containing the *E. coli* FAD synthetase. When the *Mj*FMNAT plasmid is expressed in *E. coli*, we hypothesize that if the FAD analogues are synthesized, the cells will grow to a lower extent than the *E. coli* FAD synthetase counterpart. This is because they will be deriving riboflavin from the same common pool, and these analogues (i) are unlikely to be used by the cell’s FAD-utilizing enzymes, and thus add a metabolic burden or (ii) they may act as FAD mimics and inhibit cellular functioning. The growth of three cultures - *E coli*. Bl21 (DE3) cells containing the empty vector, and the vectors with *Ec*FADS and *Mj*FMNAT – were compared. Indeed, the cells expressing *Mj*FMNAT grew ~ 0.8x that of the empty vector and *Ec*FADS controls (Figure 8A). Further, these three cultures were lysed, and the cell-free extracts were analysed on LCMS. We expect that the *Mj*FMNAT containing cells will show us a range of FAD analogues as compared to the empty vector control and the *Ec*FADS vector containing cells. FAD was detected in all the cultures, though FAD was found in relatively higher amount in cells expressing the *Ec*FADS and *Mj*FMNAT vectors as both enzymes can form FAD (Figure 8B). Additionally, *Mj*FMNAT forms other FAD analogues – FGD, FCD, FUD, and dFAD – are distinctly visible in the extracted ion chromatograms (EICs) (Figure 8B). The identity of the analogues extracted from *E. coli* were corroborated by comparing their retention time and fragmentation patterns with the FAD analogues we synthesized. Hence, we conclude that FAD nucleobase analogues can be synthesized within cellular systems using an appropriate enzyme.

**Figure 8:**
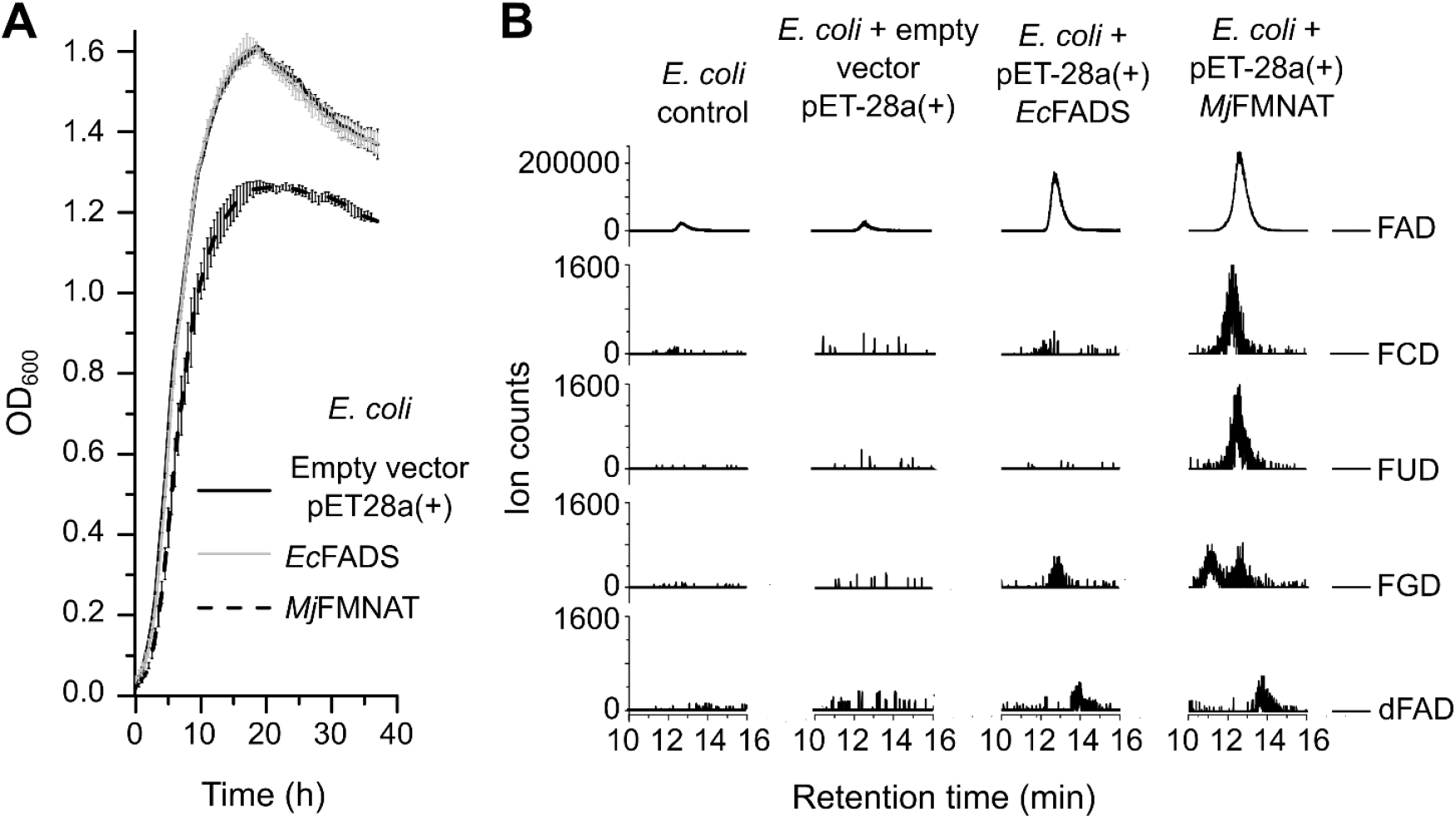
Synthesis of FAD analogues in the cell and their effect on cell growth. (A) *E. coli* with heterologously expressed *Mj*FMNAT on a plasmid has lower growth as compared to *E. coli* or *E. coli* with heterologously expressed *Ec*FADS, which show similar growth. (B) LC-MS analysis of *Mj*FMNAT expressing strain where the Extracted Ion Chromatograms (EICs) show the formation of FAD nucleoside analogues, while the *Ec*FADS and empty plasmid strains were not found to synthesize these FAD analogues.

## CONCLUSION

We set out to synthesize FAD nucleobase analogues and establish their ability to catalyse redox reactions and be produced within cells. Some routes to synthesize FAD analogues have been reported in the past but are limited by factors such as difficult-to-obtain starting materials, several steps for the synthesis, and/ or low yields. In this study, using chemical and enzymatic methods, we synthesized nucleobase analogues of FAD (FCD, FUD, FGD and dFAD) in 1-3 steps with moderate yields (in 10-51%) in 3-5 hrs. Additionally, enzymatic reactions have advantage over chemical methods as they are precise with regard to the product formed, the enzyme is required in catalytic amounts, and purification of the product can generally be achieved using green chemistry methods. Next, we showed that the FAD nucleobase analogues are viable catalysts for a representative enzyme, glutathione reductase. Finally, since the *Mj*FMNAT enzyme is promiscuous for NTPs *in vitro*, we show that its heterologous expression in *E. coli* can utilise indigenously available NTPs and FMN and can biosynthesize FAD nucleobase analogues within the cell.

The chemical and enzymatic methods we report allows us to extend the synthesis of FAD nucleobase analogues to produce artificial fluorescent FAD variants which may have several interesting applications in drug development, cell imaging, and biotechnology applications such as bioremediation^[15,44–51]^. We envision the use of these FAD nucleobase analogues as biochemical probes for cellular metabolism, and as biorthogonal catalysts to study specific steps in metabolism by engineering enzymes to utilize a FAD nucleobase analogue over the natural FAD^[16–21]^.

## MATERIALS AND METHODS

### Materials

All nucleoside triphosphates (NTPs) and 2’-deoxy nucleoside triphosphates (dNTPs) were purchased from Jena Biosciences. Chemical and reagents used in synthesis were obtained from Merck, TCI Chemicals, RANKEM chemicals, and Avra Chemicals. Genomic DNA of *Methanocaldococcus jannaschii* DSM 2661 was a gift from Biswarup Mukhopadhyay at Virginia State University. All bacterial culture, media components and reagents for biological assays were obtained from SRL chemicals, HiMedia, and TCI Chemicals. The high fidelity PrimeSTAR GXL DNA polymerase, Dpn1, and Taq polymerase used for cloning was from DSS Takara Bio USA. Plasmid and DNA purification kits were purchased from Qiagen and Agilent. All solvents used for synthesis, HPLC, and LC-MS, were obtained from RANKEM, SD Fine Chemicals and Merck. *E. coli* K-12, strain AG1 containing *Ec*GR expressing gene in pCA24N plasmid was from ASKA collection, National BioResource Project.

### Methods

#### Synthesis of 2’, 3’, 4’ -triacetyl-riboflavin 5’ -monophosphate (FMNAc)

FMN (300 mg, 0.63 mmol) was dried at high vacuum for 30 min in a round-bottomed (RB) flask. Then acetic anhydride (2 mL, 21 mmol) was added followed by 3-5 drops of ~ 55% perchloric acid (~100- 150 μL). The FMN was dissolved completely, and the solution was stirred at room temperature for 1 hr. The reaction product was precipitated out by adding diethyl ether. The mixture was filtered, and precipitate was re-dissolved in 5 mL water. First, the acetylated riboflavin was removed by workup with chloroform (10 mL x 3) and finally the product was purified by automated C18-reversed-phase chromatography using water (solvent A) and MeOH (solvent B) as mobile phase over 60 min at flow rate of 10 mL/min using the following method: 0-5 min 5% B; 10-30 min from 5% B to 50% B; 30-50 min 50% B to 100% B; 50-60 min 100% B. The chromatograms were recorded at 280 nm and 365 nm. The product was finally concentrated and dried under reduced pressure. The identity was confirmed by LC-MS and NMR (Supplementary Figure S5 and S6) and the yield obtained was 33%. The NMR spectra matches with that reported in literature^[39]^.

#### FMNAc

**^1^H NMR** (400 MHz, D_2_O) δ 7.59 (s, 1H), 7.57 (s, 1H), 5.52 (m, 2H), 5.40 (m, 1H), 3.98 - 4.23 (m, 2H), 2.51 (s, 3H), 2.34 (s, 3H), 2.26 (s, 3H), 2.23 (s, 3H), 1.69 (s, 3H). **^31^P NMR** (400 MHz, D_2_O) δ 0.59 (s). LC-MS (ESI-TOF) m/z: calculated for C_23_H_27_N_4_O_12_P [M-H]^-^ 581.1285, found 581.1286.

#### Synthesis of FNDAc

NMP (free acid, 0.15 mmol) was dried under reduced pressure for 30 min in an RB flask. All the reactions were done under nitrogen atmosphere. 400 μL of anhydrous MeCN solvent was added to the NMP followed by anhydrous N,N-dimethylaniline (60 μL, 0.5 mmol) and anhydrous triethylamine (175 μL, 1.25 mmol). The mixture was cooled and stirred in ice-water bath for 10 min. In another RB flask, TFAA (175 μL, 1.25 mmol) was dissolved in anhydrous MeCN (160 μL) and cooled in an ice-water bath. Then the TFAA solution was added to NMP mixture, and the resulted solution was stirred for 15 min at room temperature (RT). The NMP was dissolved and excess of TFAA was carefully evaporated under reduced pressure. A pale-yellow colour syrup was obtained and re-dissolved in 100 μL anhydrous MeCN followed by cooling in ice-water bath (Figure S4). In another RB flask, N-methylimidazole (30 μL, 0.4 mmol) was dissolved in 40 μL anhydrous MeCN. The solution was cooled in ice-water bath and added to trifluoroacetylated NMP solution. The mixture was stirred in ice-water bath for 15 min. In another RB flask containing 3Å molecular sieves, dry FMNAc (115 mg, 0.2 mmol, synthesized in the previous step) was dissolved in 1 mL anhydrous MeCN and anhydrous tributylamine (48 μL, 0.2 mmol) was added to it. The N-methylimidazole activated NMP solution was added to the latter mixture with stirring. The reaction was monitored on TLC (water: n-butanol: acetic acid: : 5: 12: 3). The reaction was completed in 2 hrs and quenched by addition of 10 mL of 250 mM ammonium acetate. The aqueous phase was extracted with dichloromethane (10 mL x 2) and then purified by automated C18 reversed-phase chromatography using 0.1% triethylammonium acetate in water (solvent A) and MeOH (solvent B) as mobile phase at flow rate of 10 mL/min using the following method: 0-5 min 5% B; 10-50 min from 5% B to 30% B; 50-60 min 30% B to 100% B; 60-70 min 100% B. The chromatograms were recorded at 280 nm and 365 nm. The purified product was concentrated and dried under reduced pressure and ^1^H, ^13^C, ^31^P NMR, and LC-MS were recorded (Figure S10-12). Yield obtained was 31%.

#### FCDAc

**^1^H NMR** (400 MHz, D_2_O) δ 7.96 (d, *J* = 7.7 Hz, 1H), 7.78 (s, 1H), 7.70 (s, 1H), 6.06 (d, *J* = 7.7, 1H), 5.84 (d, *J* = 3.7 Hz, 1H), 5.55 – 5.57 (m, 2H), 5.45 (br, 1H), 5.16 (br, 1H), 4.97 (br, 1H), 4.32 – 4.14 (m, 7H), 2.58 (s, 3H), 2.43 (s, 3H), 2.27 (s, 3H), 2.25 (s, 3H), 1.72 (s, 3H). **^13^C NMR** (400 MHz, D2O) δ 172.85, 172.53, 165.08, 160.87, 157.25, 157.22, 156.29, 150.58, 149.94, 141.59, 139.26, 134.25, 134.09, 130.91, 130.77, 116.21, 96.14, 89.16, 82.56 (d, *J*_CP_ = 33.4 Hz), 74.30, 70.75 (d, *J*_CP_ = 21.8 Hz), 70.44, 69.84, 69.05, 64.42 (d, *J*_CP_ = 17.4 Hz), 63.78 (d, *J*_CP_ = 17.5), 45.05, 20.81, 20.63, 20.25, 19.70, 18.56. **^31^P NMR** (400 MHz, D_2_O) δ −10.94 (s). LC-MS (ESI-TOF) m/z: calculated for C_32_H_39_N_7_O_19_P_2_ [M-H]^-^ 886.1698, found 886.1699.

#### Deacetylation of FNDAc

The entire reaction was performed under nitrogen atmosphere. ~10 mg of dry FNDAc was dissolved in 1 mL anhydrous MeOH as a solvent and the reaction was initiated by the addition of 100 μL of 0.5 M NaOMe in MeOH. In ~5 min, the product was precipitated out as FND is not soluble in MeOH. The reaction was quenched just after precipitation with 100 μL of 0.5 M HCl. The product was purified by automated C18 reversed-phase chromatography using 0.1% triethylammonium acetate in water (solvent A) and MeOH (solvent B) as mobile phase at flow rate of 10 mL/min using the following method: 0-5 min 5% B; 10-50 min from 5% B to 20% B; 50-70 min 20% B to 100% B; 70-80 min 100% B. The chromatograms were recorded at 280 nm and 365 nm. The purified product was concentrated and dried under reduced pressure and ^1^H, ^13^C, and ^31^P NMR and LC-MS were recorded (Figure S13-15). Yield for FCD was 17% with respect to CMP as starting material.

#### FCD

**^1^H NMR** (400 MHz, D_2_O) δ 7.98 (d, *J* = 7.7 Hz, 1H), 7.79 (s, 1H), 7.63 (s, 1H), 6.07 (d, *J* = 7.7 Hz, 1H), 5.79 (d, *J* = 4.0 Hz, 1H), 5.05 (dd, *J* = 12.8, 10.4 Hz, 1H), 4.68 (dd, *J* = 12.8, 1.2 Hz, 1H), 4.44 – 4.37 (m, 1H), 4.31 – 4.08 (m, 8H), 4.00 (dd, *J* = 7.2, 4.9 Hz, 1H), 2.50 (s, 3H), 2.38 (s, 3H). **^13^C NMR** (600 MHz, D_2_O) δ 160.70, 159.19, 156.09, 151.20, 149.10, 148.24, 140.86, 137.82, 132.65, 132.20, 129.99, 128.65, 115.38, 94.05, 87.67, 81.16 (d, *J*_CP_ = 37.1 Hz), 72.76, 70.86, 69.54 (d, *J*_CP_ = 31.0 Hz), 67.52, 67.32, 65.56 (d, *J*_CP_ = 22.0 Hz), 62.70 (d, *J*_CP_ = 20.7 Hz), 45.91, 19.15, 17.02.**^31^P NMR** (400 MHz, D_2_O) δ −10.62 (d, *J*_PP_ = 49.5 Hz,), −11.39 (d, *J*_PP_ = 49.5 Hz). LC-MS (ESI-TOF) m/z: calculated for C_26_H_33_N_7_O_16_P_2_ [M-H]^-^ 760.1381, found 760.1376.

### Activity assay of *Mj*FMNAT

We tested the activity of *Mj*FMNAT with the various NTPs – ATP, GTP, CTP, UTP, and dATP. For this reaction, 170 μM FMN, 10 mM NTP, 7 mM MgCl_2_, 14 mM DTT, 50 mM Tris-HCl pH 8.0 and 5 μM enzyme were mixed in a total volume of 500 μL and heated at 70°C for 12 hrs. After 12 hrs, the reaction was quenched by adding 60 mM acetic acid (final concentration). The formation of FAD and its analogues were analysed on the TLC and reversed-phase HPLC. The FAD and FAD analogues formed in the *Mj*FMNAT reaction were purified and collected using the same HPLC method. For the collection of analogues 200 μL of samples were injected through the autosampler (50 μL at a time) and collected manually. The solvents from the samples were evaporated using Centrivap DNA concentrator (Labconco corporation). The dried powder of analogues was stored at −20°C. To determine the yield of dFAD and FGD, the dried powder was dissolved in 200 μL of buffer and injected into the HPLC. The concentrations of dFAD and FGD were determined from area under the curve, assuming these have same absorption as FAD. The percentage yield of the product was calculated with respect to the initial concentration of the substrate used in the reaction. The final yields of the enzymatically synthesized FGD and dFAD were 51% and 45%, respectively.

LC-MS (ESI-TOF) m/z: calculated for FGD (C_27_H_33_N_9_O_16_P_2_) [M-H]^-^ 800.1442, found 800.1425.

LC-MS (ESI-TOF) m/z: calculated for dFAD (C_27_H_33_N_9_O_14_P_2_) [M-H]^-^ 768.1544, found 768.1545.

### *In vivo* formation of FAD analogues

in 50 mL LB broth, secondary culture was started by inoculation with 0.5 mL primary culture of BL21 (DE3) cells containing pET28a, pET28a-*Ec*FADS, or pET28a-*Mj*FMNAT construct resistant to kanamycin and only culture of BL21 (DE3) was grown without kanamycin at 37°C. After 18 hrs, 50 mL of culture was palleted down by centrifugation at 4°C (5500 rpm x 15 min). The pellets were resuspended in 2 mL of MeOH and sonicated at 1 sec pulse on and 1 sec pulse off for 3 minutes at 100% amplitude. The lysate was centrifuged at 4°C (5500 rpm x 15 min) and the supernatant was filtered with 0.22-micron filters. The supernatant then concentrated on high-vac and used for LC-MS analysis.

### Activity assay of *E. coli* glutathione reductase (*Ec*GR) with FAD and its analogues

To analyze the reduction capability of the FAD nucleobase analogues, we used the *Ec*GR enzyme. Briefly, this assay involves the reduction of FAD to FADH_2_ with NADPH which gets oxidized to NADP and then FADH2 reduces the disulfide bridge of the enzyme which further reduces GSSG to GSH^[52]^. We assayed *Ec*GR by monitoring decrease in absorbance of NADPH in reaction mixture at 340 nm in a 96 well plate where NADPH is oxidized to NADP by GSSG^[41]^. The reaction mixture contained 0.1M potassium phosphate buffer pH 7, 1 mM EDTA, 1 mM GSSG, and 0.1 mM NADPH of 200 μL final volume. Reaction was started by adding 0.1 μM (final concentration) of reconstituted enzyme at 30°C.

### TLC, HPLC, and LC-MS

All reactions were analysed on silica gel TLC and visualized under ultraviolet light (UV) at 365 nm using mobile phase n-butanol: acetic acid: water in the ratio 12:3:5.

HPLC for analysing the syntheses as well as the enzymatic reactions was conducted with a UV-Vis diode array detector equipped Agilent 1260 Infinity-II series high-performance liquid chromatography (HPLC) system. For all reactions, a Phenomenex Gemini with dimensions 250 x 4.6 mm, 5μm NX-C18 column was used. The solvents used for the HPLC were 10 mM ammonium acetate buffer pH 6.0 (solvent A) and MeOH (solvent B). A 60 min gradient method with a flow rate of 0.5 ml/min was employed, beginning with 5% B at 0 min, holding at 5% B until 15 min, then a gradient to 80% B until 40 min, and then a gradient to 5% B until 55 min, and holding at 5% B until 60 min.

ScieX X500R-QTOF Mass spectrometer system attached to Exion-LC series UHPLC was used to perform all LC-MS analyses. The analytes were separated using the same method and column as in HPLC method. Negative ESI mode was used for all analyses with −4500 V ion spray voltage, medium de-clustering potential (−80V) and low collision energy (5V) at 400°C. ScieX-OS Explorer was used to acquire and process the data and to obtain total ion chromatogram (TIC), extracted ion chromatogram (EIC), and the mass spectra of desired molecules.

## ACKNOWLEDGMENTS

We are grateful to Biswarup Mukhopadhyay at Virginia State University for a gift of the genomic DNA of *Methanocaldococcus jannaschii* DSM 2661 that we have used for cloning. We thank Pulak Ghosh from the Srivatsan lab, Jayashree Rajput from the Hotha lab, and members of the Chakrapani and the Khan labs at IISER Pune for help with organic synthesis and use of their equipment. We thank Gayathri Pananghat and S. G. Srivatsan for insightful discussions. We acknowledge the help received from the LC-MS and NMR facilities and their managers Saddam Shekh and Nitin Dalvi at IISER Pune.

