## Supplementary Information for "Efficient chemical and enzymatic syntheses of FAD nucleobase analogues and their analysis as enzyme cofactors"

#### SUPPLEMENTARY METHODS

##### Alternate method for synthesis of FCD

FMN (300 mg, 0.63 mmol) was dried at high vacuum for 30 min in a round bottom (RB) flask. Then acetic anhydride (2 mL, 21 mmol) was added followed by 3-5 drops of ~ 55% perchloric acid (~100 µL). The solution was stirred at room temperature for 1 hr. The reaction product was precipitated out by adding diethyl ether to remove acetic acid and anhydride. The mixture was filtered, and the precipitate was re-dissolved in 5 mL water. The aqueous phase was extracted with chloroform (10 mL x 3) to remove acetylated riboflavin. Mixture of acetylated FMN was obtained which was used directly for further coupling reaction. CMP (free acid, 50 mg, 0.15 mmol) was dried under reduced pressure for 30 min in an RB flask. All the reactions were done in the nitrogen atmosphere. 400 µL anhydrous MeCN was added to the CMP followed by anhydrous N, N-dimethylaniline (60 µL, 0.5 mmol) and anhydrous triethylamine (175 µL, 1.25 mmol). The mixture was cooled and stirred in ice-water bath for 10 min. In another RB flask, trifluoro acetic anhydride (TFAA) (175 µL, 1.25 mmol) was dissolved in anhydrous 160 µL MeCN and cooled in ice-water bath. Then the TFAA solution was added to CMP mixture, and the solution was stirred for 15 min at room temperature (RT). The access of TFAA was carefully evaporated under reduced pressure. A pale-yellow colour syrup was obtained and re-dissolved in 100 µL anhydrous MeCN followed by cooling in ice-water bath. In another RB flask, N-methylimidazole (30 µL, 0.4 mmol) was dissolved in 40 µL anhydrous MeCN. The solution was cooled in ice-water bath and added to trifluoroacetylated CMP solution. The mixture was stirred in ice-water bath for 15 min. In another RB flask containing 3Å molecular sieves, dry mixture of acetylated FMN (3 times excess of CMP) was dissolved in anhydrous MeCN. The N-methylimidazole activated NMP solution was added to latter mixture and the reaction was kept stirring for 2 hrs. The reaction was monitored on TLC (water: n-butanol: acetic acid: : 5: 12: 3). The reaction was quenched by addition of 10 mL of 250 mM ammonium acetate. The aqueous phase was extracted with dichloromethane (10 mL x 2) and dried under reduced pressure. It was directly deacetylated with 50 mM NaOMe in MeOH for 3-5 min where FCD and FMN precipitated in MeOH. However, some of FCD breaks to cyclic FMN and CMP in this condition same as FAD[1]. FCD was purified by C18 reversed phase chromatography using 10mM ammonium acetate in water (solvent A) and MeOH (solvent B) as mobile phase using same method as mentioned in main paper. The purified product was concentrated and dried under reduced pressure (Yield 13%).

### Supplementary information

#### Synthesis of FUD

UMP (free acid, 50mg, 0.15 mmol) was dried under reduced pressure for 30 min in an RB flask. All the reactions were done in the nitrogen atmosphere. 400 µL anhydrous MeCN was added to the UMP followed by anhydrous N, N-dimethylaniline (60 µL, 0.5 mmol) and anhydrous triethylamine (350 µL, 2.5 mmol). The mixture was cooled and stirred in ice-water bath for 10 min. In another RB flask, TFAA (175 µL, 1.25 mmol) was dissolved in 160 µL anhydrous MeCN and cooled in ice-water bath. Then the TFAA solution was added to UMP mixture and the solution was stirred for 15 min at RT. The UMP was dissolved and access of TFAA was carefully evaporated under reduced pressure. A pale-yellow colour syrup was obtained and re-dissolved in 100 µL anhydrous MeCN followed by cooling in ice-water bath. In another RB flask, N-methylimidazole (30 µL, 0.4 mmol) was dissolved in 40 µL anhydrous MeCN. The solution was cooled in ice-water bath and added to trifluoroacetylated UMP solution. The mixture was stirred in ice-water bath for 15 min. In another RB flask containing 3Å molecular sieves, dry 2', 3', 4'-triacetyl-riboflavin 5' -monophosphate (115 mg, 0.2 mmol) was dissolved in 1 mL anhydrous MeCN and anhydrous tributylamine (500 µL, 2 mmol) was added to it. The N-methylimidazole activated UMP solution was added to later mixture and the reaction was kept stirring. The reaction was monitored on TLC (water: n-butanol: acetic acid: : 2.5: 6: 1.5). The reaction was completed in 2 hrs and quenched by addition of 10 mL of 250 mM ammonium acetate. The aqueous phase was extracted with dichloromethane (10 mL x 2) and then purified by automated C18 reversed phase chromatography using 0.1% triethylammonium acetate in water (solvent A) and MeOH (solvent B) as mobile phase at flow rate of 10 mL/min using the following method: 0-5 min 5% B; 10-50 min from 5% B to 30% B; 50-60 min 30% B to 100% B; 60-70 min 100% B. The chromatograms were recorded at 280 nm and 365 nm. The purified product was concentrated and dried under reduced pressure, and $^{1}H$, $^{13}C$, and $^{31}P$ NMR and LC-MS were recorded (Figure S16-S18). Yield was 11%.

**$^{1}$H NMR** (600 MHz, $D_2O$) δ 7.84 (s, 1H), 7.82 (s, 1H), 7.72 (s, 1H), 5.85 – 5.78 (m, 2H), 5.11 – 5.02 (m, 1H), 4.44 – 4.37 (m, 1H), 4.32 – 4.08 (m, 8H), 4.04 – 3.96 (m, 1H), 2.52 (s, 3H), 2.40 (s, 3H). **$^{13}$C NMR** (600 MHz, $D_2O$) δ 164.02, 159.57, 156.10, 149.61, 148.84, 148.28, 139.50, 137.56, 132.85, 132.41, 130.03, 128.62, 115.11, 100.57, 86.48, 81.27, 72.02, 70.75, 69.27, 67.69, 65.39, 65.38, 63.02, 45.81, 18.97, 16.81. **$^{31}$P NMR** (600 MHz, $D_2O$) δ -8.07 (br), -8.75 (br). LC-MS (ESI-TOF) m/z: calculated for $C_{26}H_{32}N_6O_{17}P_2$ $[M-H]^-$ 761.1226, found 761.1228.

### Supplementary information

#### Restriction free cloning of *Mj*FMNAT

The FMN adenylyl transferase gene (*MJ*1179) from *Methanocaldococcus jannaschii* was cloned into pET28a vector between the restriction sites *NdeI* and *BamHI* using restriction free cloning[2]. The MJ1179 gene was amplified by polymerase chain reaction using the following primers: Forward 5'-ATGAAAAAGAGGGTAGTTACCGC-3' and Reverse TTAGATTTTAATCTCTTTATTGCAG. The complementary sites for the *NdeI* and *BamHI* restriction sites were amplified at the ends of the PCR amplified gene by a second PCR using forward primer GTGCCGCGCGGCAGCCATATGAAAAAGAGGGTAGTTACCGC and reverse primer CGACGGAGCTCGAATTCGGATCCTTAGATTTTAATCTCTTTATTGCAG. This gene sequence with complementary flanking regions at terminal ends is called the megaprimer. This megaprimer was used to amplify the pET28a vector with the gene inserted in a third round of PCR. The parent plasmid was digested with Dpn1 (NEB), and the vector containing *MJ*1179 was transformed into *E. coli* DH5α chemically competent cells for plasmid isolation.

#### Purification of *Mj*FMNAT

The purified plasmid containing the *MJ*1179 gene corresponding to the enzyme *Mj*FMNAT was transformed into *E. Coli* BL21 (DE3) chemically competent cells. The cells were grown in Luria-bertani broth at 37°C. When O.D. of the culture was 0.6, it was induced with 1 mM IPTG at 30°C. After the post-induction time of 8 hrs, cells were harvested and lysed in lysis buffer (100 mM Tris-HCl pH 8.0 and, 300 mM NaCl, 0.025% beta-mercaptoethanol, 0.1 mM PMSF). Lysed cells were centrifuged (18000 rpm x 30 min), and the supernatant was loaded into pre-equilibrated Ni-NTA column (100 mM Tris-HCl pH 8.0 and, 300 mM NaCl, 0.025% beta-mercaptoethanol) using FPLC system (Akta pure, GE healthcare). The protein was eluted with 250 mM imidazole buffer (Tris-HCl pH 8.0 and, 300 mM NaCl, 0.025% beta-mercaptoethanol, 250 mM imidazole). For activity assays, concentration was measured using $A_{280}$ having predicted molar extinction coefficient 5960 $M^{-1}$ $cm^{-1}$.

#### Purification of *E. coli* glutathione reductase (*Ec*GR)

*E. Coli* K12 AG1 cells transformed with the *Ec*GR in a pCA24n vector, were grown in LB broth at 37°C. When O.D. of the culture reached to 0.5-0.6, it was induced with 0.5 mM IPTG at 28°C. After the post-induction time of 8 hrs, cells were harvested and lysed in lysis buffer (100 mM Tris-HCl pH 8.0 and, 300 mM NaCl, 0.025% beta-mercaptoethanol, 0.1 mM PMSF). Lysed cells were centrifuged (18000 rpm x 30 min), and the supernatant was loaded into pre-equilibrated Ni-NTA column (100 mM Tris-

HCl pH 8.0 and, 300 mM NaCl, 0.025% beta-mercaptoethanol). The protein was eluted with 250 mM imidazole buffer (Tris-HCl pH 8.0 and, 300 mM NaCl, 0.025% beta-mercaptoethanol, 250 mM imidazole).

To obtain the apoprotein, saturated ammonium sulphate solution was used[3]. 1 mL KBr (3 M solution) was added to 1 ml protein in an ice-cold centrifuge tube followed by precipitation by dropwise addition of 1 mL saturated ammonium sulfate, pH 2.2, under gentle swirling. After 1 min, additional 4 ml of saturated ammonium sulfate was added. The solution was centrifuged (18000 rpm x 30 min) and the precipitate was redissolved in 1 ml of 0.1 M phosphate buffer, pH 7 to give the apoprotein and the residual denatured protein was removed by centrifugation. The apoprotein was again reconstituted with FAD or FAD analogues by incubating it with an excess of the cofactor at 25°C for 15 min. Then, the excess cofactor was removed by passing the protein through desalting column equilibrated in 0.1 M phosphate buffer, pH 7. For activity assays, concentration was measured using Bradford assay.

**RMSD of FAD molecules in FAD-binding proteins**

We wished to obtain a quantitative comparison of conformations of FAD in FAD-binding enzymes. Root Mean Square Deviation (RMSD) is the square root of the average of squared errors and is represented as RMSD= $[(\sum(x_e-x_0)^2)/n]^{0.5}$ (where $x_e$=expected value, $x_o$=observed value and $n$=total number of values)[4]. RMSD of atomic positions is a quantitative measure of the average distance between the corresponding atoms of two superimposed molecules (FAD in this case). 24 FAD-binding protein structures deposited in the PDB-Protein Data Bank (https://www.rcsb.org/) were used for this purpose[5]. The "match command" on UCSF Chimera was used to perform the best fit (superposition) of specified atoms by moving the atoms of one FAD molecule onto the second FAD molecule followed by calculation of the RMSD values[6]. 53 atoms present in an FAD molecule (all except the hydrogen atoms) were considered for these calculations. Pairwise RMSD calculations were done for 24 such FAD molecules. The FAD molecule bound to Thioredoxin disulfide reductase in *Campylobacter jejuni* (PDB ID: 3R9U) was arbitrarily chosen as the reference molecule with 0.0 Å, and the RMSD values for each of the other 23 enzyme-bound FADs with respect to the reference molecule were plotted (Figure 1B).

### Supplementary information

**SUPPLEMENTARY FIGURES**

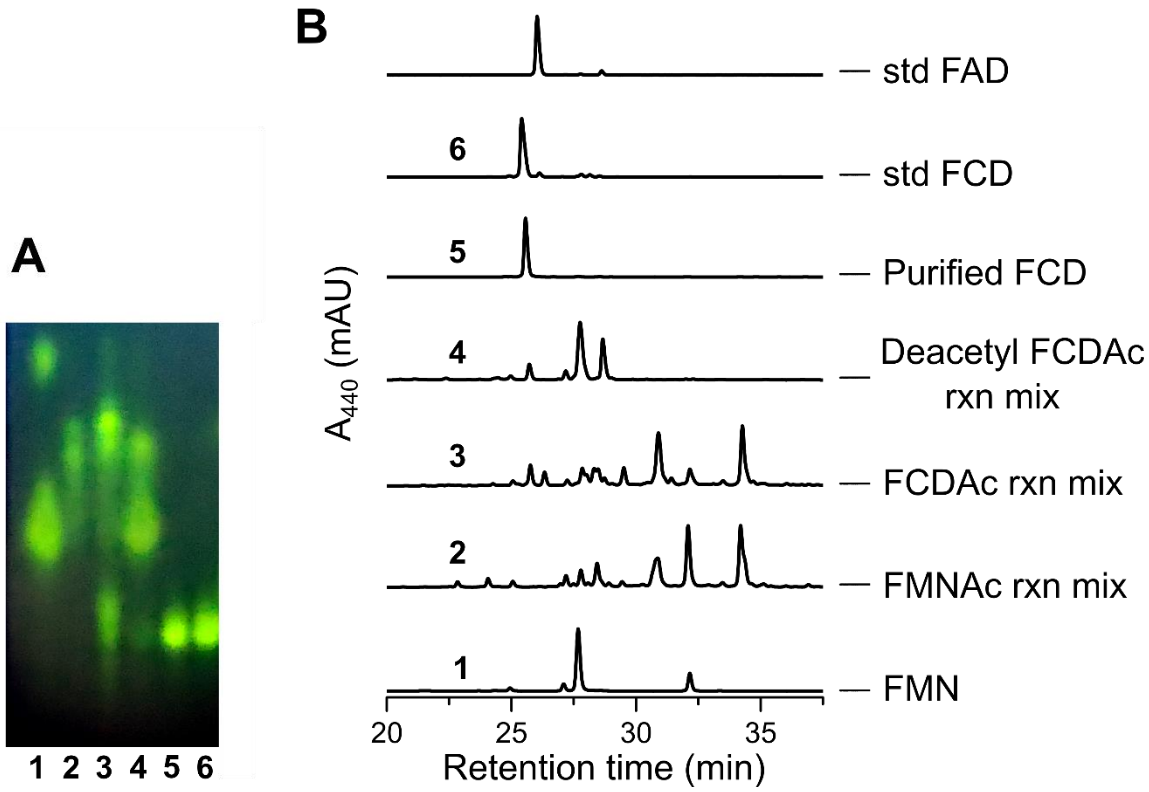


**Figure S1: Synthesis of FCD from FMN via FMNAc and FCDAc. (A)** Each step of synthesis was followed on TLC where Lanes are as 1- FMN with RF contamination (top), 2- FMNAc reaction mixture, 3- FCDAc reaction mixture, 4- deacetylation of FCDAc reaction mixture, 5- purified FCD, and 6- standard FCD. (B) HPLC chromatogram at each step where FMNAc reaction mixture shows multiple peaks for mixture of acetylated FMN. Similarly, FCDAc also shows the peaks for acetylation at different stages which on deacetylation gives final product of FCD with degradation product (cyclic FMN) and FMN.

### Supplementary information

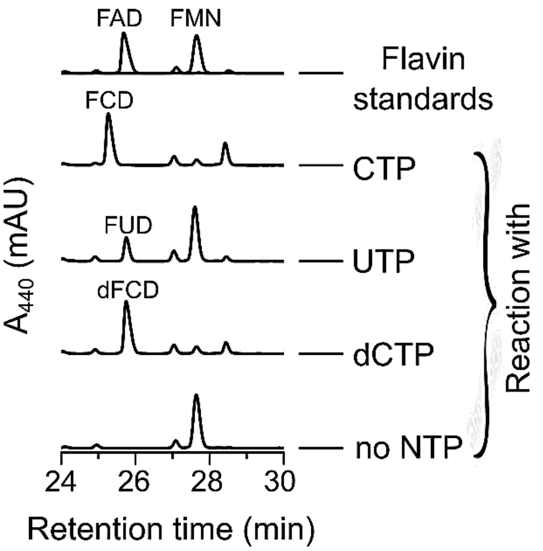


**Figure S2. Characterisation of *Mj*FMNAT reaction with NTPs and dCTP.** HPLC chromatogram shows the conversion of FMN to FNDs and dFCD with different NTPs by MjFMNAT enzyme (25.5-26 min).

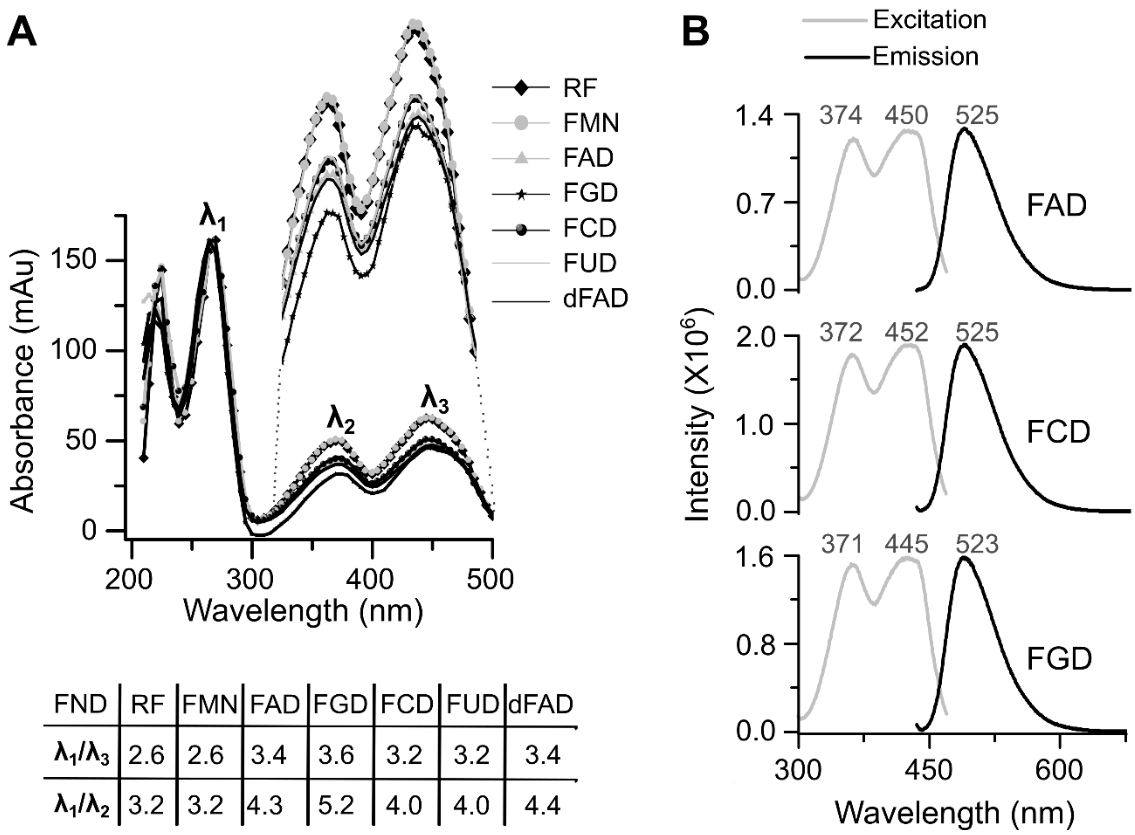


| FND | RF | FMN | FAD | FGD | FCD | FUD | dFAD |
|---|---|---|---|---|---|---|---|
| $\lambda_1/\lambda_3$ | 2.6 | 2.6 | 3.4 | 3.6 | 3.2 | 3.2 | 3.4 |
| $\lambda_1/\lambda_2$ | 3.2 | 3.2 | 4.3 | 5.2 | 4.0 | 4.0 | 4.4 |

**Figure S3. Spectrophotometry of FAD analogues.** (A) UV-visible spectra of RF, FMN, FAD, and FAD analogues. Each molecule shows different $\lambda_1/\lambda_2$ ($\lambda_{270}$: $\lambda_{370}$) and $\lambda_1/\lambda_3$ ($\lambda_{270}$: $\lambda_{450}$) depending on the nucleobase as shown in the table. RF and FMN overlaps with no nucleobase, FAD and dFAD overlaps with adenine base, similarly FCD and FUD has same ratio with the pyridine bases, and FGD shows the highest ratio with guanine base. (B) comparison of Fluorescence spectra of pyridine and purine nucleobase analogues of FAD.

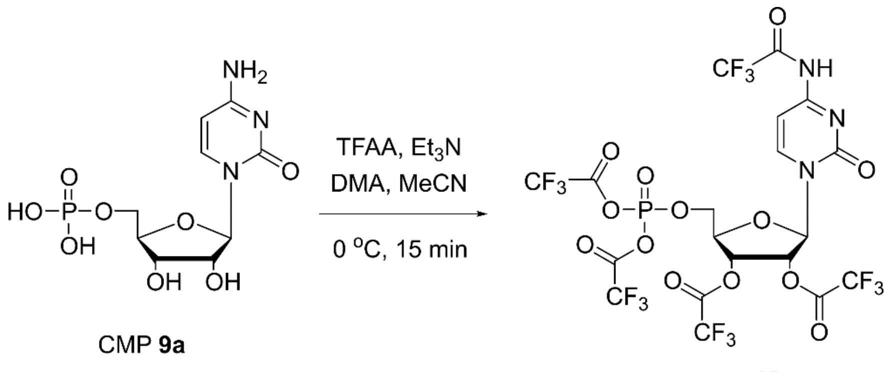


**Figure S4: Activation of phosphate group of CMP with TFAA.**

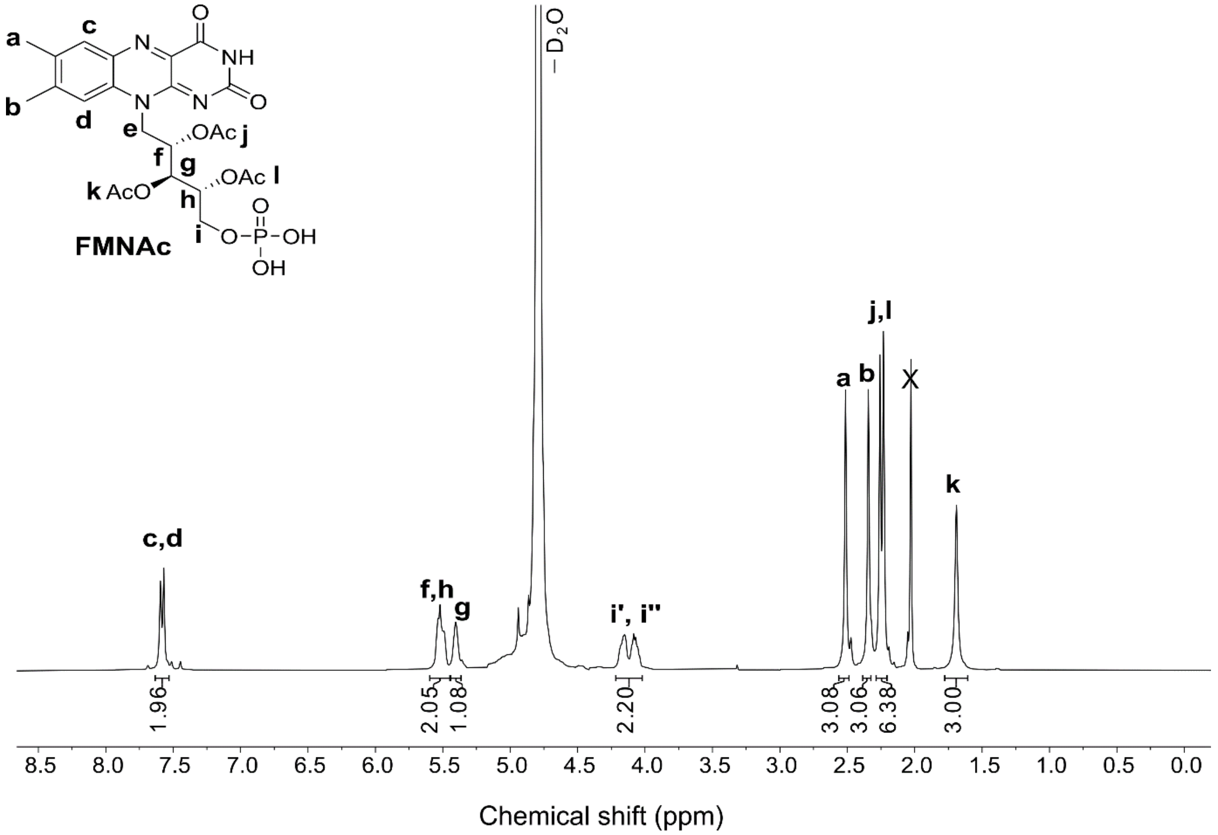


**Figure S5. $^{1}H$ NMR (400 MHz, $D_2O$) of FMNAc (8)**

### Supplementary information

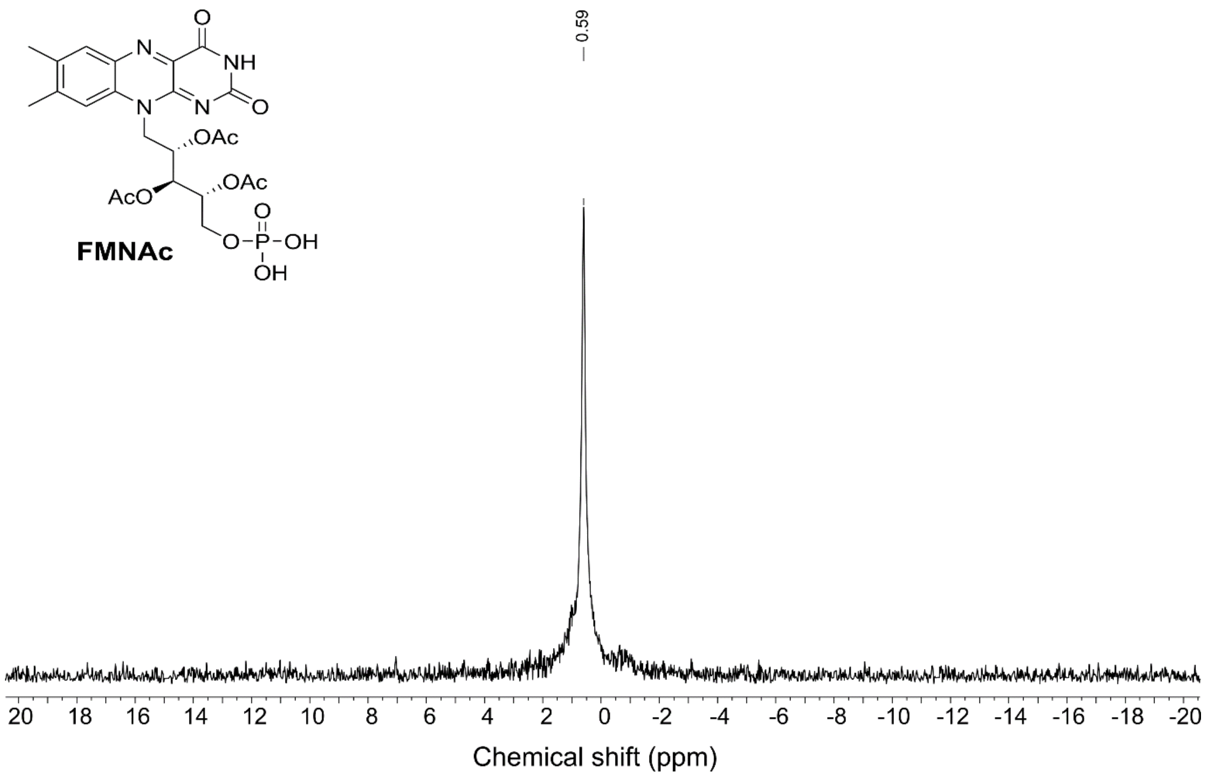


**Figure S6. $^{31}$P NMR (400 MHz, $D_2O$) of FMNAc (8)**

### Supplementary information

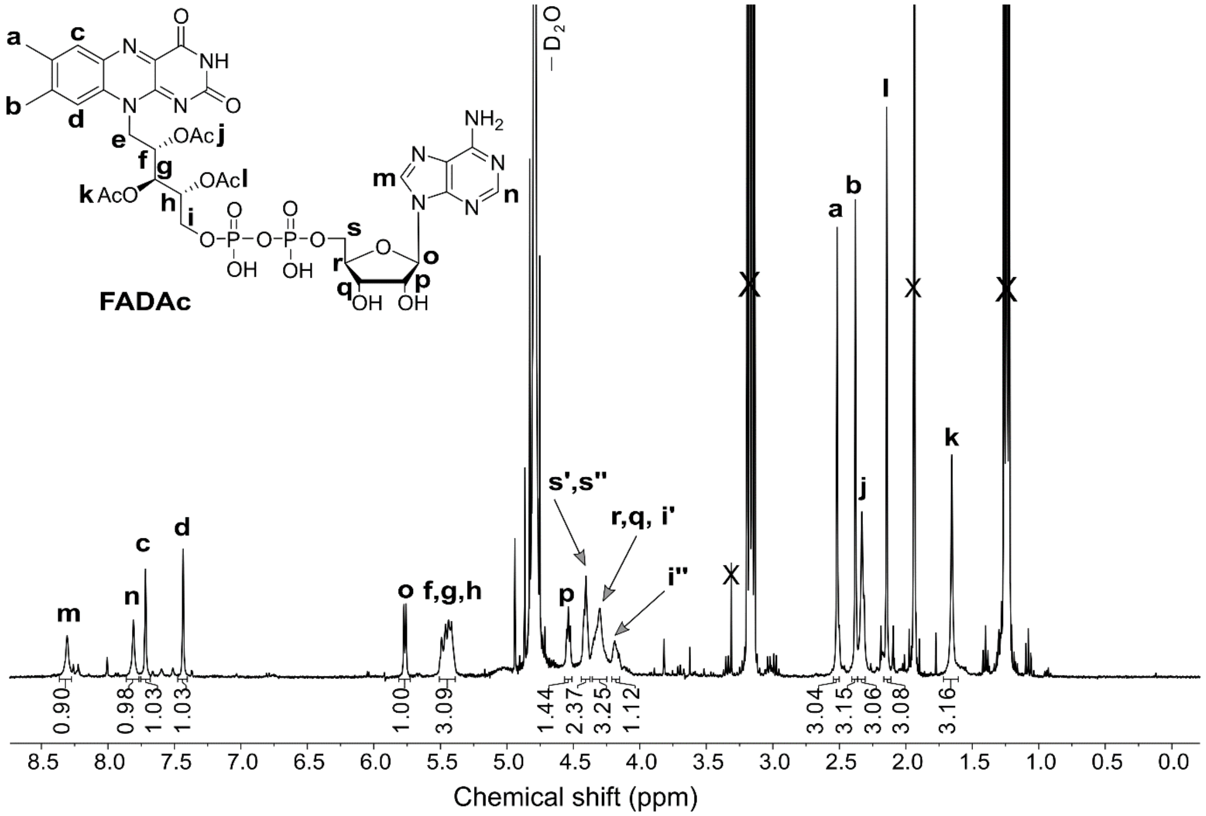


**Figure S7. $^{1}$H NMR (400 MHz, $D_2O$) of FADAc**

### Supplementary information

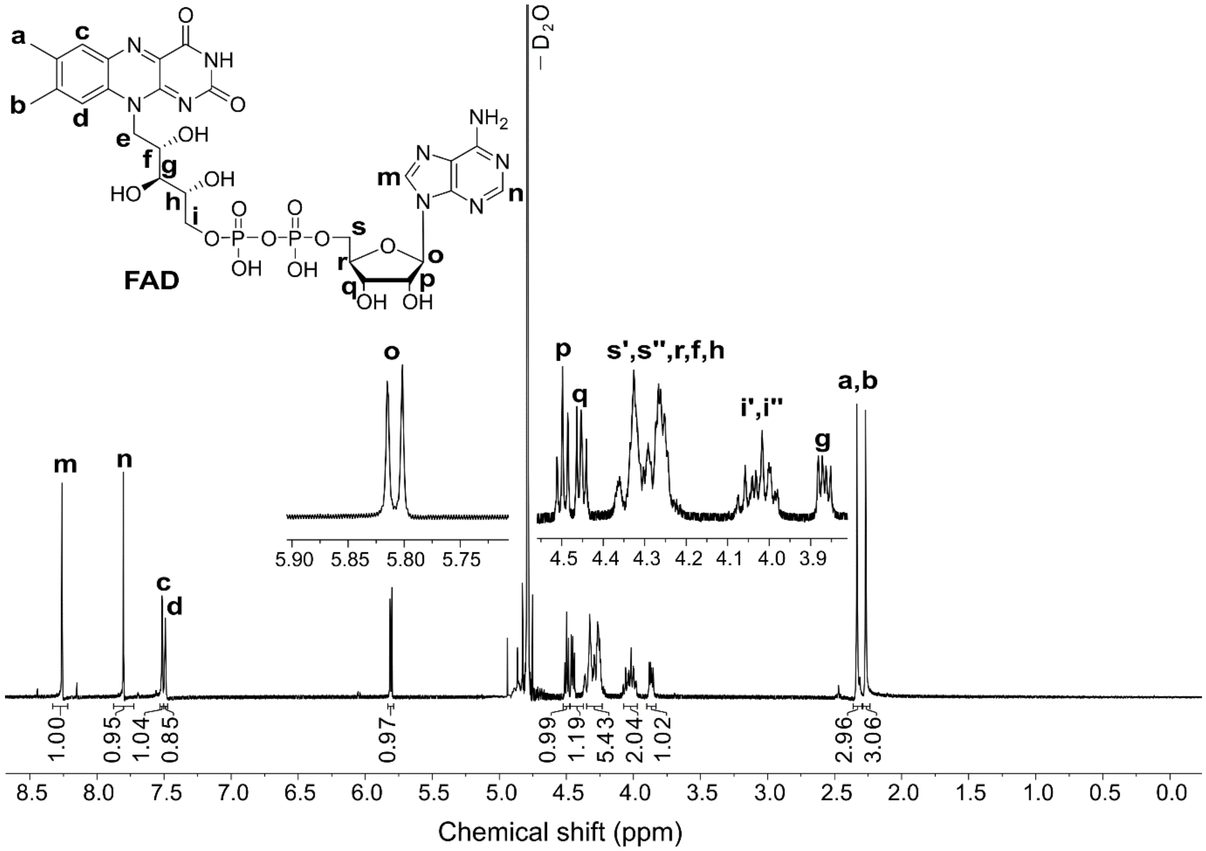


**Figure S8. $^1$H NMR (600 MHz, $D_2O$) of FAD (3)**

### Supplementary information

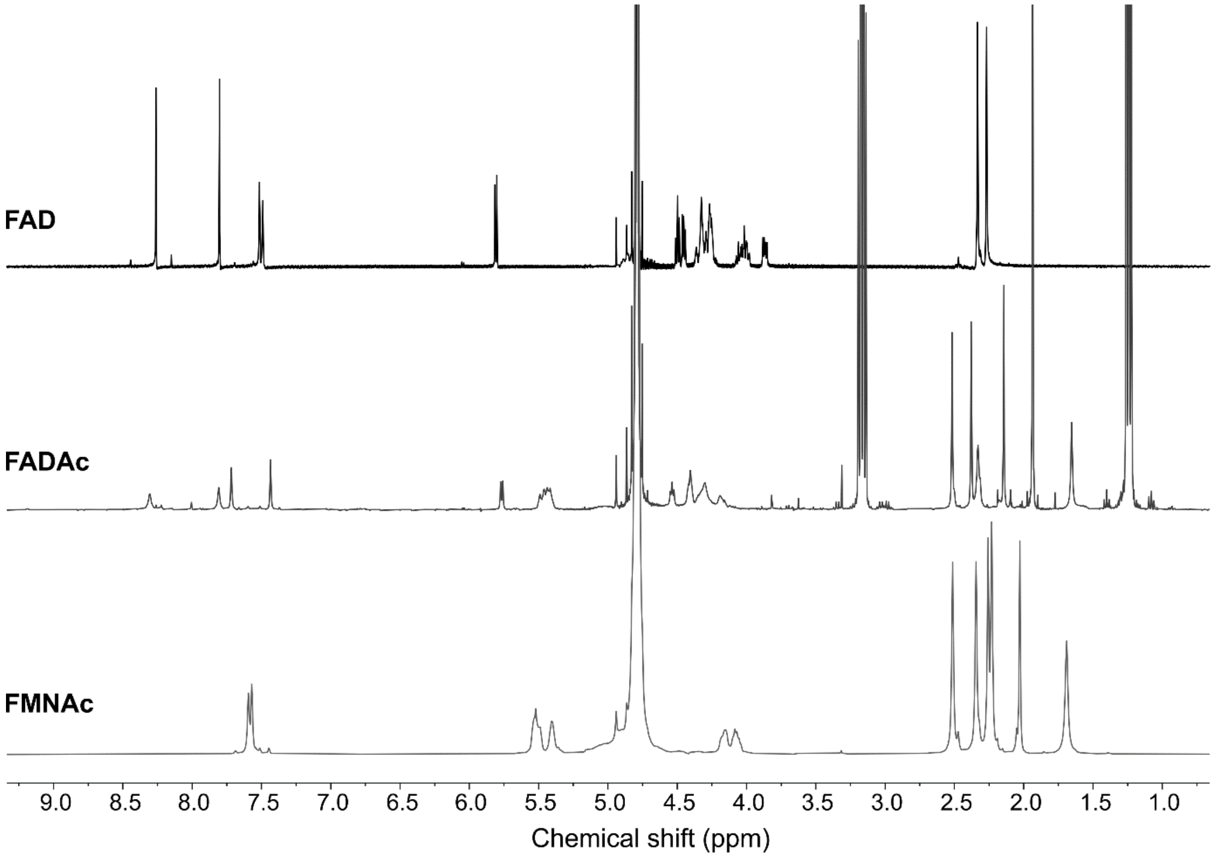


**Figure S9. Comparison of NMR spectra of intermediates (FMNAc and FADAc) and the final product FAD.**

### Supplementary information

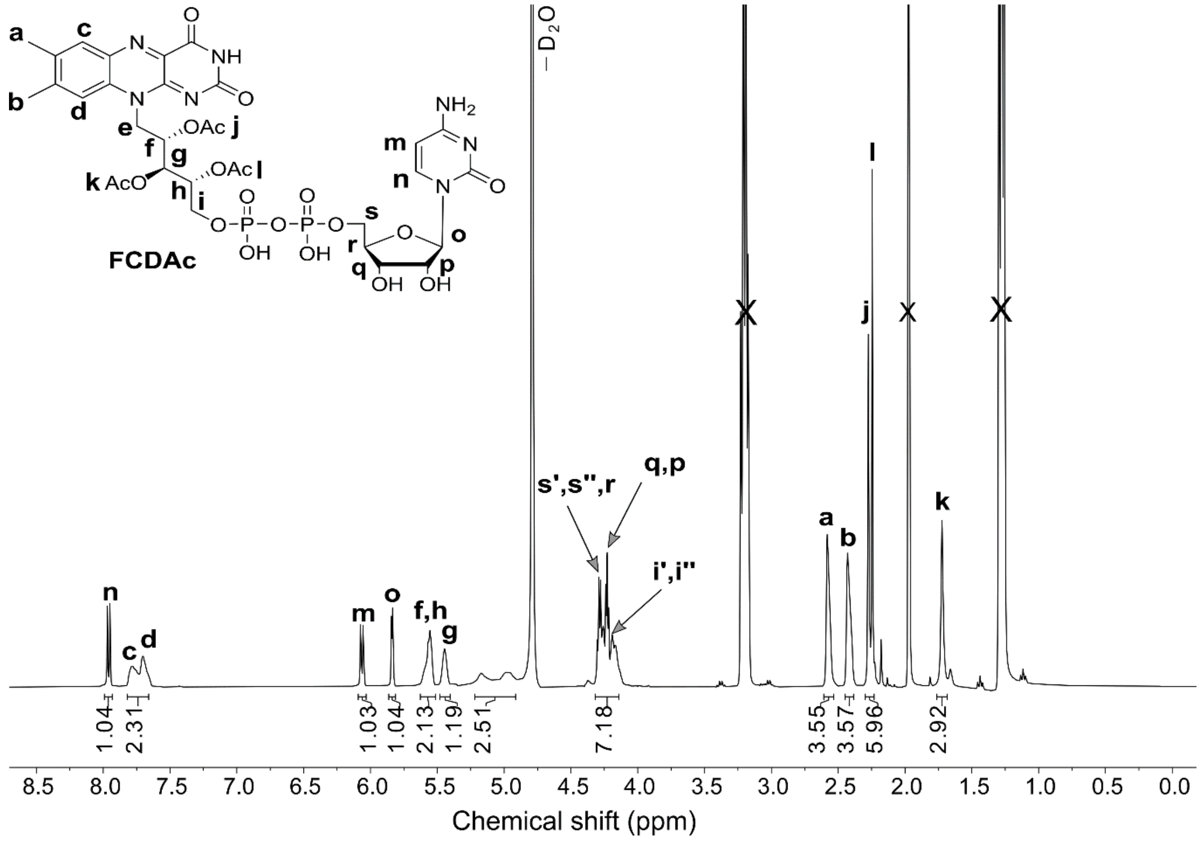


**Figure S10. $^{1}$H NMR (400 MHz, $D_2O$) of FCDAc (11)**

### Supplementary information

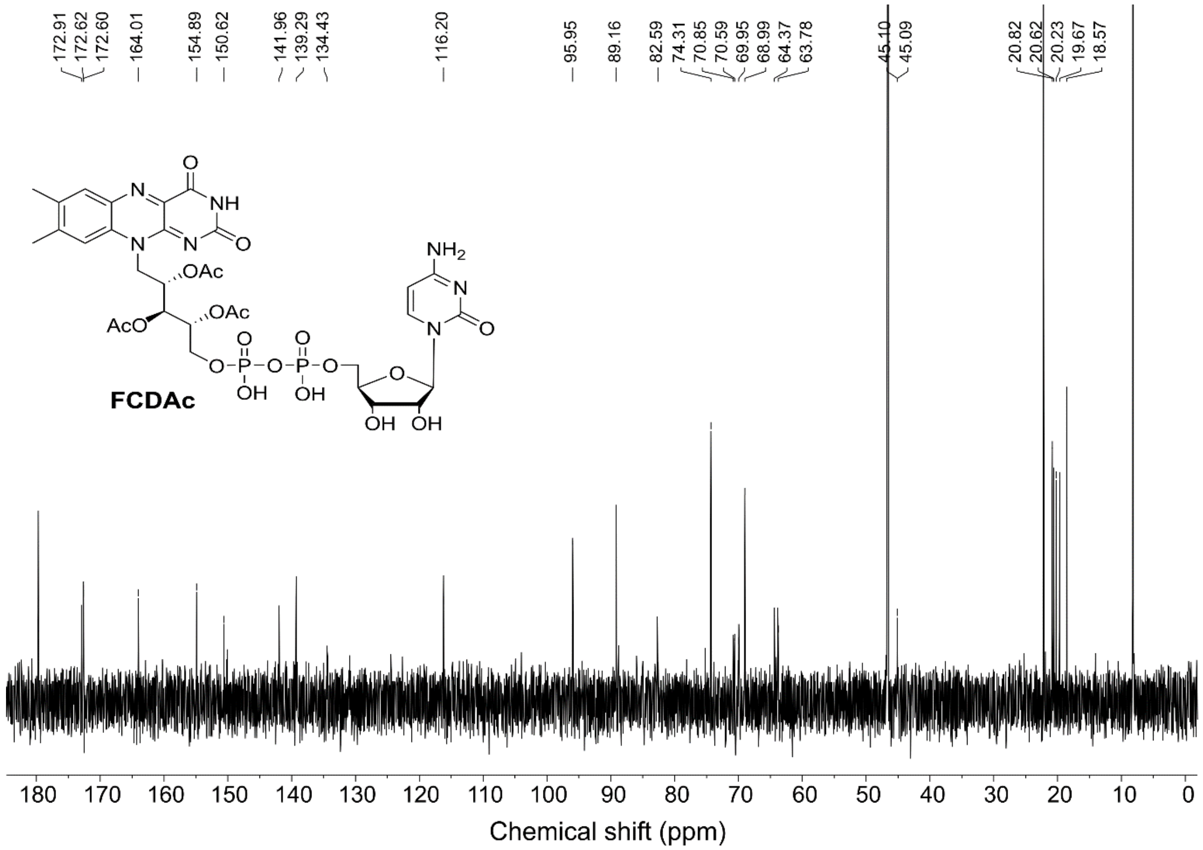


**Figure S11. $^{13}$C NMR (400 MHz, $D_2O$) of FCDAc (11)**

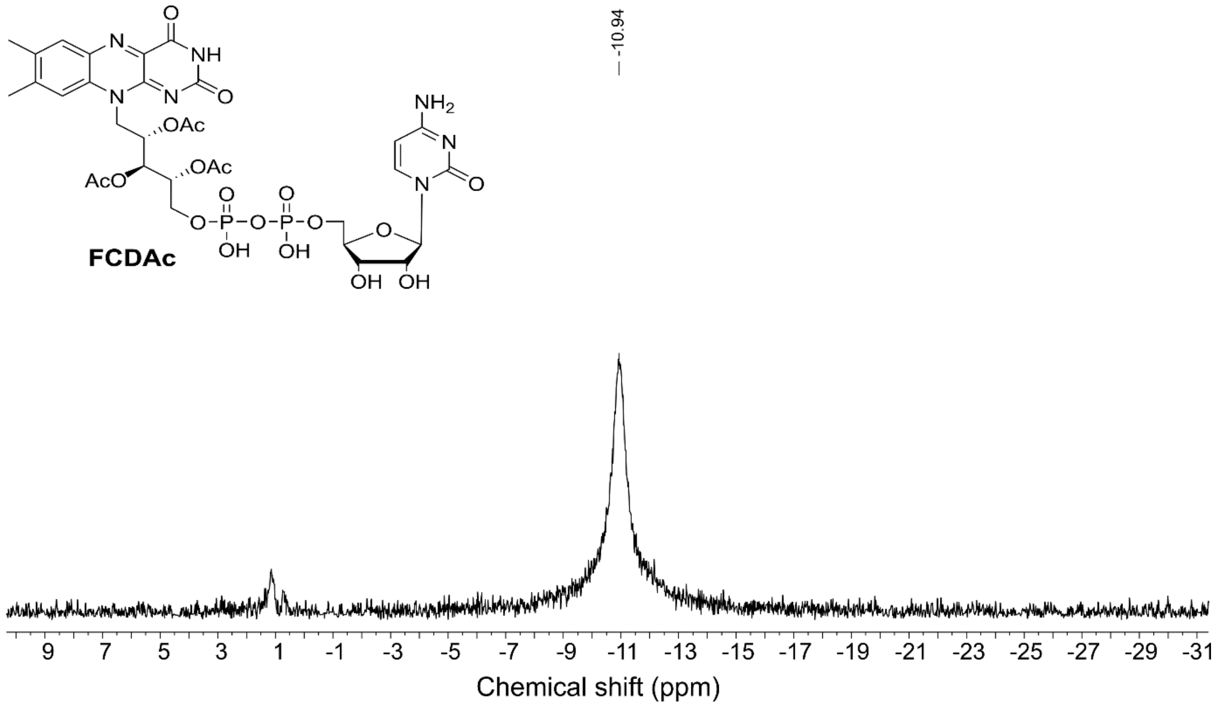


**Figure S12. $^{31}$P NMR (400 MHz, $D_2O$) of FCDAc (11)**

### Supplementary information

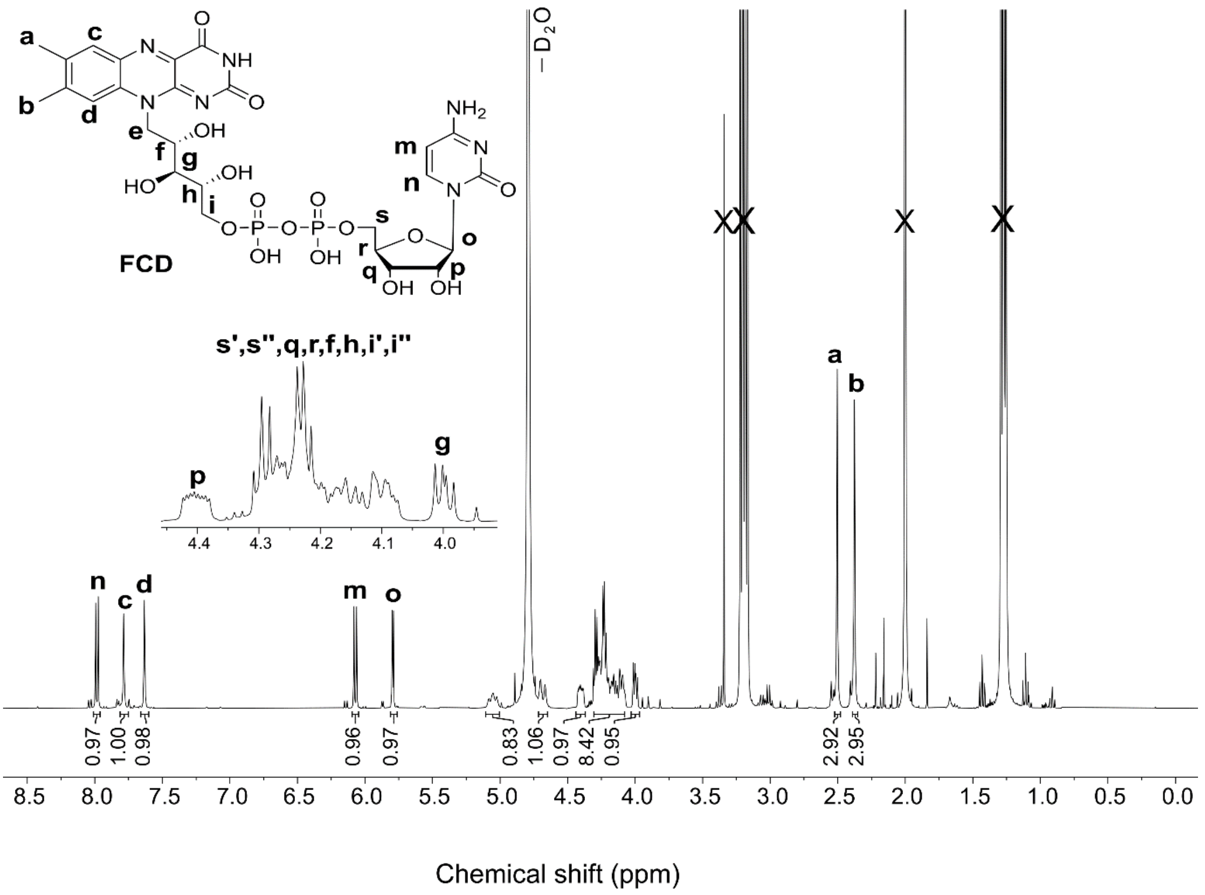


**Figure S13. $^{1}$H NMR (400 MHz, $D_2O$) of FCD (5)**

### Supplementary information

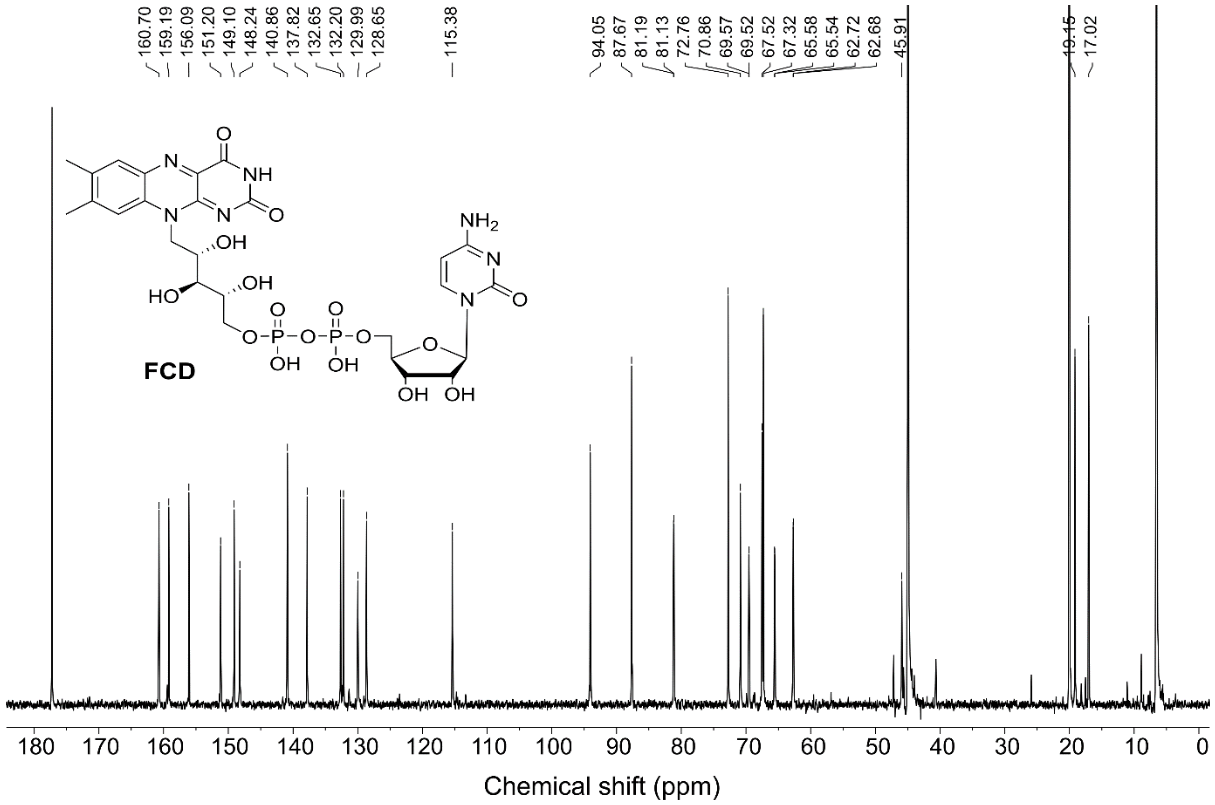


**Figure S14.** **$^{13}C$ NMR (400 MHz, $D_2O$) of FCD (5)**

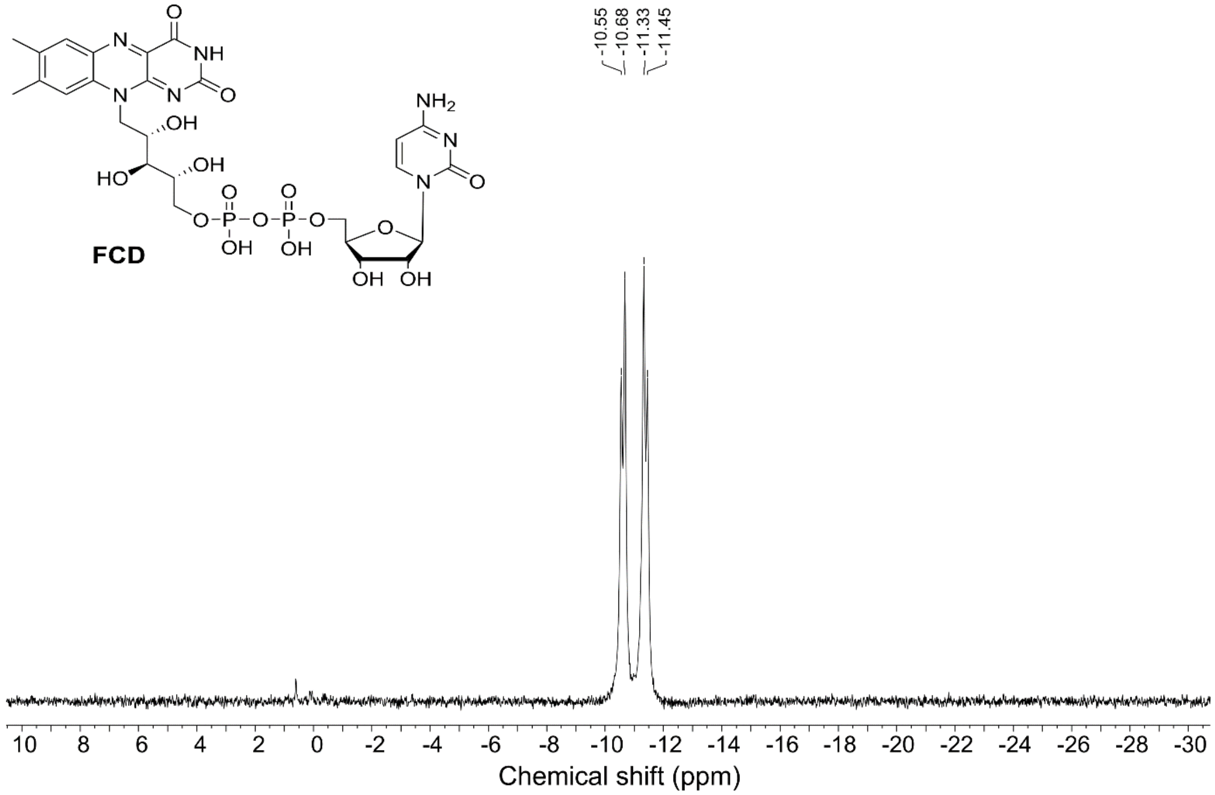


**Figure S15.** **$^{31}P$ NMR (400 MHz, $D_2O$) of FCD (5)**

### Supplementary information

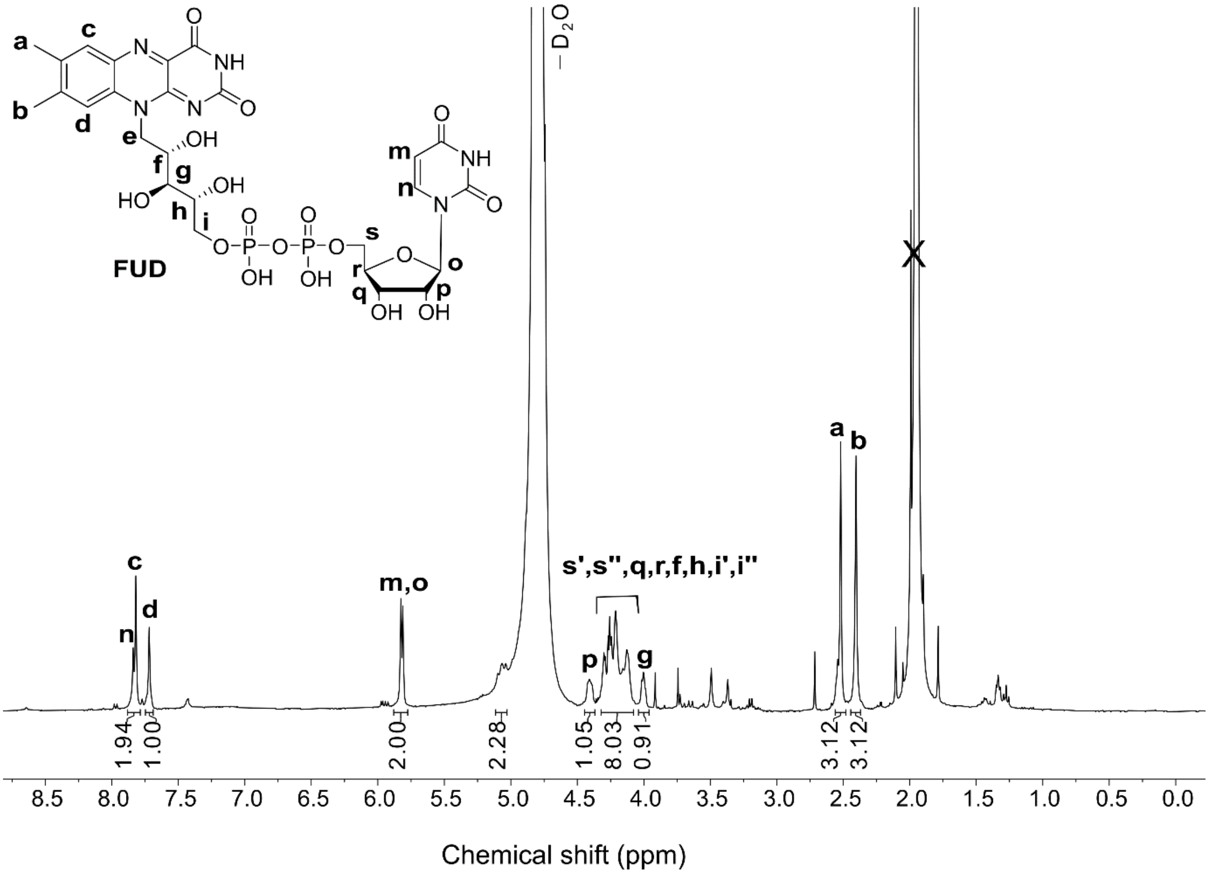


**Figure S16.** $^{1}$**H NMR (400 MHz,** $D_2O$**) of FUD (6)**

### Supplementary information

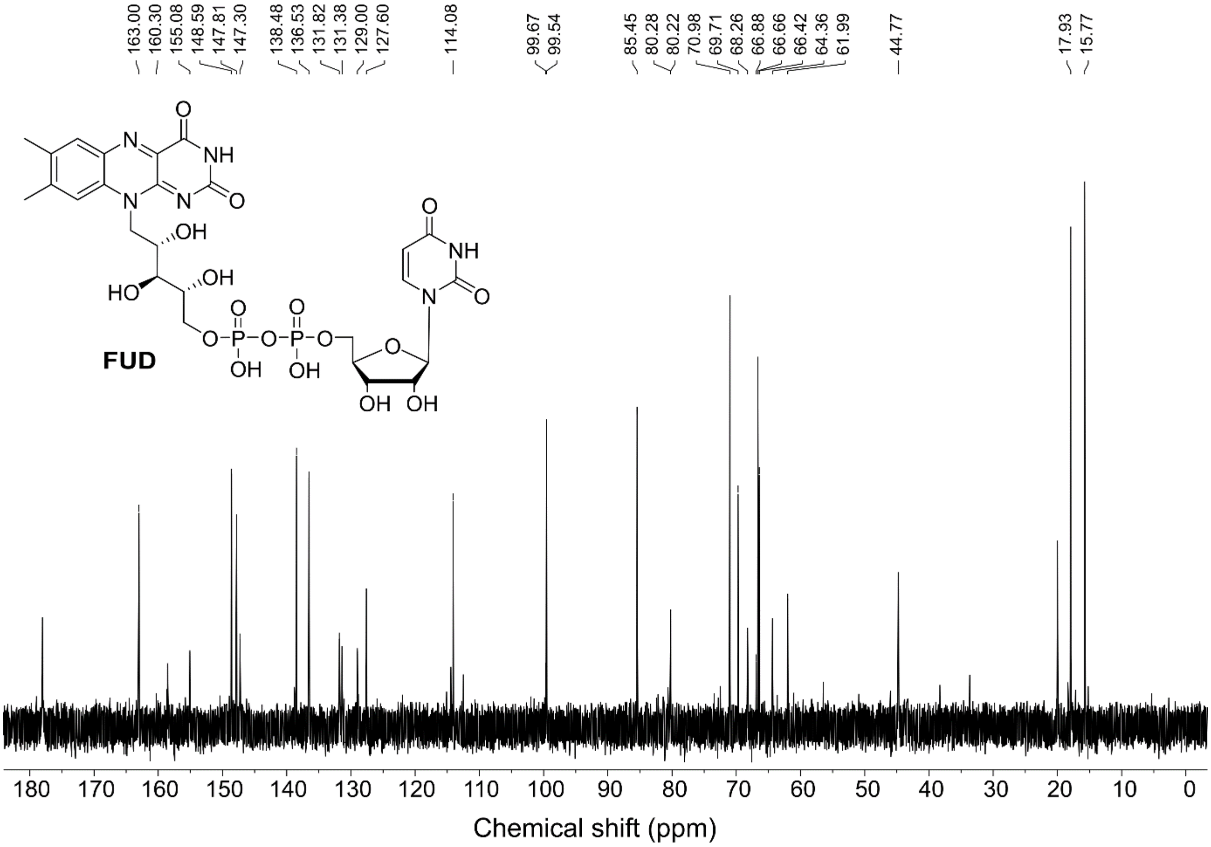


**Figure S17. $^{13}$C NMR (400 MHz, $D_2O$) of FUD (6)**

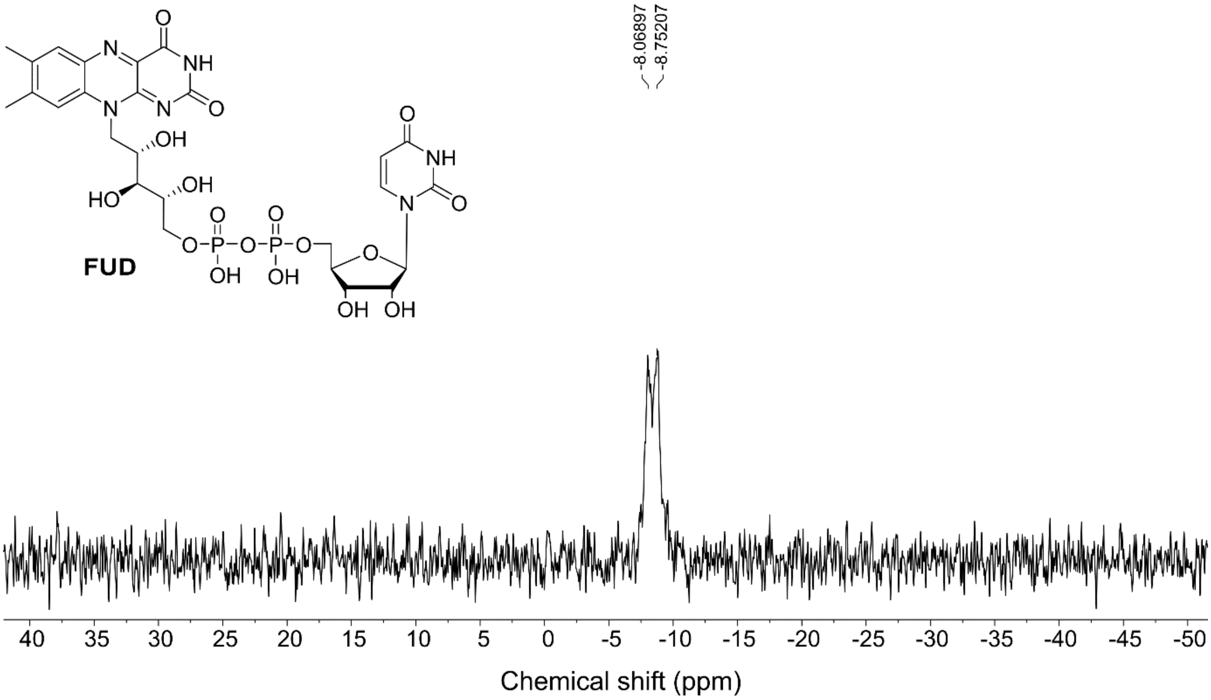


**Figure S18. $^{31}$P NMR (400 MHz, $D_2O$) of FUD (6)**

### Supplementary information

| S. No. | Method | description | Molecule | Ref. | yield | time | Steps |
|---|---|---|---|---|---|---|---|
| 1 | Chemical synthesis | Condensation of 2’: 3‘-O-isoPropylideneadenosine-5’ Benzyl Phosphorochloridate and monothallous riboflavin-5’ phosphate in phenol followed by removal of thallous chloride with dry ether. Deprotection in acid | FAD | CHRISTIE, KENNER, and TOD. *J. Chem. Soc.* (1954) | 6.6% | 10 h | 4 |
| 2 | Chemical synthesis | Coupling of FMN and AMP using di-p-tolyl carbodiimide in pyridine | FAD<br>FID | Huennekens and Kilgour. *JACS* (1955) | <4% | 24 h | 1 |
| 3 | Large scale chemical synthesis | Direct condensation of FMN and NMP in presence of trifluoroaceticacid anhydride | FAD<br>FGD<br>FCD<br>FUD | DeLuca and Kaplan. *JBC* (1956) | <1% | 16 h | 1 |
| | | | FCD | Mashhadi et al. *Biochemistry* (2010) | Not reported | 18 h | 1 |
| 4 | Chemical synthesis | Coupling of adenosine-5' phosphoramidate or phosphoromorpholidate and riboflavin-5' phosphate in a mixture of pyridine and o-chlorophenol | FAD | Moffatt and Khorana. *JACS* (1958) | 40% | 4 d | 2 |
| | | | FdeAD<br>FGD, FdeGD<br>FCD, FdeCD<br>FTD<br>FHD, FdeHD<br>FXD, FdeXD | McCormick, Chassy and Tsibris. *Biochimica et Biophysica Acta* (1964) | - | - | 2 |
| | | | FdeAD<br>FGD, FdeGD<br>FCD, FdeCD<br>FTD<br>FHD, FdeHD<br>FXD, FdeXD | Tsibris, McCormick, and Wright. *Biochemistry* (1965) | - | - | 2 |

### Supplementary information

| | | | | | | | |
|---|---|---|---|---|---|---|---|
| | | | Flavin-8-bromoadenine dinucleotide | McCormick and Opar. *Journal of Medicinal Chemistry* (1969) | 28% | 7 d | 2 |
| 5 | Chemical synthesis | Mono (tri-n-octylammonium) salt of Adenosine-5' phosphate coupled with Diphenyl phosphochloridate DMF and dioxan followed by attack of mono (tri-n-octylammonium) flavin mononucleotide in DMF and pyridine. | FAD | Michelson. *Biochimica et Biophysica Acta* (1964) | 70-80% | 5 h | 2 |
| | | | 5-deazaflavin adenine dinucleotide | Smit. *Recueil des Travaux Chimiques des Pays-Bas* (1984) | - | 12-15 h | 3 |
| | | | Azido-FAD | Koberstein. *European Journal of Biochemistry* (1976) | 25% | 5 h | 2 |
| | | | $N^6$-(6-Carboxyhexyl)-FAD | Stocker, Hecht, And Buckmann. *European Journal of Biochemistry* (1996) | 30% | 12-17 h | 4 |
| 6 | Chemical synthesis | Coupling of nucleoside S'-phosphorothioate with ammonium salt of FMN in dry pyridine | FAD | Hata and Nakagawa. *JACS* (1970) | 51% | 5 h | 1 |
| 7 | Enzymatic synthesis | Coupling of FMN and NTP using *Ca*FADS | - | Spencer, Fisher, and Walsh. *Biochemistry* (1976) | - | 4-18 h | 1 |

**Table S1:** Other reported methods for synthesis of FAD and its nucleoside analogues.

### Supplementary information

| | Enzyme concentration (μM) | Cofactor concentration (μM) | Cofactor binding (%) |
|---|---|---|---|
| Purified *Ec*GR | 199.7 | 195.96 | 98.13 |
| apo *Ec*GR | 86.97 | 5.34 | 6.14 |
| *Ec*GR + FAD | 93.25 | 83.44 | 89.47 |
| *Ec*GR + FCD | 98.23 | 83.91 | 85.43 |
| *Ec*GR + FUD | 82.72 | 74.9 | 90.55 |

**Table S2:** Cofactor binding of purified, apo, and reconstituted *E. coli* glutathione reductase (*Ec*GR).

| S. No. | PDB ID | Name of Enzyme | Organism Name | RMSD (wrt PDB: 3R9U) (Å) |
|---|---|---|---|---|
| 1 | 1EGC | Medium Chain Acyl CoA dehydrogenase | *Human* | 2.974 |
| 2 | 2G37 | L-Proline dehydrogenase | *Thermus thermophilus* | 3.401 |
| 3 | 3R9U | Thioredoxin disulfide reductase | *Campylobacter jejuni* | 0 |
| 4 | 6BZ0 | Dihydrolipoyl dehydrogenase | *Acinetobacter baumannii* | 0.801 |
| 5 | 2VFR | Alditol Oxidase | *Streptomyces coelicolor* | 3.267 |
| 6 | 4I58 | Cyclohexylamine Oxidase | *Microbacterium oxydans* | 0.388 |
| 7 | 1TZL | Pyranose-2-oxidase | *Peniophora sp* | 0.77 |
| 8 | 1F0X | D-lactate dehydrogenase | *E. coli* | 3.036 |
| 9 | 1OJD | Monoamine Oxidase B | *Human* | 0.488 |
| 10 | 2BVH | 6-hydroxy-D-nicotine oxidase | *Paenarthrobacter nicotinovorans* | 3.584 |
| 11 | 7PBG | 4-ethyl phenol oxidase | *Gulosibacter chungangensis* | 3.08 |
| 12 | 6PXS | iminodiacetate oxidase | *Chelativorans sp. BNC1* | 0.927 |
| 13 | 3PQB | GilR | *Streptomyces griseoflavus* | 3.342 |
| 14 | 1E8G | Vanillyl Alcohol Oxidase | *Penicillium simplicissimum* | 3.056 |
| 15 | 1W1O | Cytokinin Dehydrogenase | *Zea mays* | 3.477 |
| 16 | 4E0H | Sulfhydryl oxidase erv1 | *Saccharomyces cerevisiae S288C* | 3.505 |
| 17 | 4DQK | Cytochrome P450 BM3 | *Priestia megaterium* | 3.019 |
| 18 | 6ECI | MSMEG_5243 | *Mycolicibacterium smegmatis MC2 155* | 4.295 |
| 19 | 5EZ7 | oxidoreductase PA4991 | *Pseudomonas aeruginosa PAO1* | 0.816 |
| 20 | 2I0Z | FAD utilizing dehydrogenase | *Bacillus cereus* | 0.368 |

| | | | | |
|---|---|---|---|---|
| 21 | 1TVC | methane monooxygenase reductase | *Methylococcus capsulatus* | 3.907 |
| 22 | 1JR8 | Sulfhydryl oxidase Erv2p | *Saccharomyces cerevisiae* | 3.515 |
| 23 | 2WSI | FAD Synthetase | *Saccharomyces cerevisiae* | 4.31 |
| 24 | 2OAL | Tryptophan halogenase-RebH | *Lentzea aerocolonigenes* | 1.324 |

**Table S3:** List of 24 FAD-binding enzymes and their RMSD values.
